# Efficient pedigree recording for fast population genetics simulation

**DOI:** 10.1101/248500

**Authors:** Jerome Kelleher, Kevin R. Thornton, Jaime Ashanderf, Peter L. Ralph

**Affiliations:** Big Data Institute, University of Oxford; Ecology and Evolutionary Biology, University of California at Irvine; Ecology and Evolutionary Biology, University of California at Los Angeles; Institute for Ecology and Evolution, University of Oregon

## Abstract

In this paper we describe how to efficiently record the entire genetic history of a population in forwards-time, individual-based population genetics simulations with arbitrary breeding models, population structure and demography. This approach dramatically reduces the computational burden of tracking individual genomes by allowing us to simulate only those loci that may affect reproduction (those having non-neutral variants). The genetic history of the population is recorded as a succinct tree sequence as introduced in the software package `msprime`, on which neutral mutations can be quickly placed afterwards. Recording the results of each breeding event requires storage that grows linearly with time, but there is a great deal of redundancy in this information. We solve this storage problem by providing an algorithm to quickly ‘simplify’ a tree sequence by removing this irrelevant history for a given set of genomes. By periodically simplifying the history with respect to the extant population, we show that the total storage space required is modest and overall large efficiency gains can be made over classical forward-time simulations. We implement a general-purpose framework for recording and simplifying genealogical data, which can be used to make simulations of any population model more efficient. We modify two popular forwards-time simulation frameworks to use this new approach and observe efficiency gains in large, whole-genome simulations of one to two orders of magnitude. In addition to speed, our method for recording pedigrees has several advantages: (1) All marginal genealogies of the simulated individuals are recorded, rather than just genotypes. (2) A population of N individuals with M polymorphic sites can be stored in *O*(*N* log *N* + *M*) space, making it feasible to store a simulation’s entire final generation as well as its history. (3) A simulation can easily be initialized with a more efficient coalescent simulation of deep history. The software for recording and processing tree sequences is named tskit.

**Author Summary:** Sexually reproducing organisms are related to the others in their species by the complex web of parent-offspring relationships that constitute the pedigree. In this paper, we describe a way to record all of these relationships, as well as how genetic material is passed down through the pedigree, during a forwards-time population genetic simulation. To make effective use of this information, we describe both efficient storage methods for this embellished pedigree as well as a way to remove all information that is irrelevant to the genetic history of a given set of individuals, which dramatically reduces the required amount of storage space. Storing this information allows us to produce whole-genome sequence from simulations of large populations in which we have not explicitly recorded new genomic mutations; we find that this results in computational run times of up to 50 times faster than simulations forced to explicitly carry along that information.

## Introduction

Since the 1980’s, coalescent theory has enabled computer simulation of the results of population genetics models identical to that which would be produced by large, randomly mating populations over long periods of time without actually requiring simulation of so many generations or meioses. Coalescent theory thus had three transformative effects on population genetics: first, giving researchers better conceptual tools to describe *gene trees* and thus bringing within-population trees into better focus; second, producing analytical methods to estimate parameters of interest from genetic data; and finally, providing a computationally feasible method to produce computer simulations of population genetics processes. However, these powerful advances came with substantial caveats: the backwards-in-time processes that are described by coalescent theory are only *Markovian*, and thus feasible to work with, under the following important assumptions: (a) random mating, (b) neutrality, (c) large population size, and (d) small sample size relative to the population size. The first two assumptions can be side-stepped to a limited extent (Hudson 1990; Neuhauser and Krone 1997), but it remains a challenge to map results of coalescent models onto species that are distributed across continuous geographical space (Barton et al. 2010; Kelleher et al. 2014) and/or have large numbers of loci under various sorts of selection. Usually, the relationship between the life history of a species – fecundity and mortality schedules, density-dependent effects on fitness, and demographic fluctuations – are all absorbed into a single compound parameter, the coalescence rate. More mechanistic models are possible using “forwards–backwards” simulations, that first simulate population size changes forwards in time and then thread a coalescent backwards (Ray et al. 2010), but these still require the assumptions above to be met for each subpopulation. The last assumption is no longer safe, either - for example, a recent study (Martin et al. 2017) simulated 600,000 samples of human chromosome 20 to examine biases in GWAS. Several studies have now shown that in samples approaching the size of the population, genealogical properties may be distorted relative to the coalescent expectation (Bhaskar et al. 2014; Maruvka et al. 2011; Wakeley and Takahashi 2003). These considerations, and increasing computational power, have led to a resurgence of interest in large forwards-time, individual-based simulations. For instance, Harris and Nielsen (2016) used SLiM (Messer 2013) to simulate ten thousand human exomes to assess the impact of genetic load and Neanderthal introgression on human genetic diversity. Sanjak et al. (2017) used fwdpp (Thornton 2014) to simulate a series of models of quantitative traits under mutation-selection balance with population sizes of 2 × 10^4^ diploids in stable populations and populations growing up to around 5 × 10^5^ individuals, using the output to explore the relationship between the genotype/phenotype model and GWAS outcomes.

Modern computing power easily allows simulations of birth, death and reproduction in a population having even hundreds of millions of individuals. However, if our interest lies in the resulting genetic patterns of variation - and often, the point of such simulations is to compare to real data - then such simulations must record each individual’s genome. As samples of most species’ genomes harbor tens or hundreds of millions of variant sites, carrying full genotypes for even modest numbers of individuals through a simulation can quickly become prohibitive. To make matters worse, a population of size N must be simulated across many multiples of N generations to produce stable genetic patterns (Wakeley 2005; Wright 1931). Because of this computational burden, even the fastest simulation frameworks such as SLiM 2 (Haller and Messer 2017) and fwdpp (Thornton 2014) can “only” simulate tens of megabases of sequence in tens of thousands of individuals for tens of thousands of generations. In practice, current state-of-the-art simulation software may take on the order of weeks to simulate models of large genomic regions without selection (Hernandez and Uricchio 2015; Thornton 2014), and existing simulation engines differ in how efficiently they calculate fitnesses in models with selection (Thornton 2014). These population and region sizes are still substantially short of whole genomes (hundreds to thousands of megabases) for many biological population sizes of interest.

However, it is thought that most genetic variation is selectively neutral (or nearly so). By definition, neutral alleles carried by individuals in a population do not affect the population process. For this reason, if one records the entire genealogical history of a population over the course of a simulation, simply laying down neutral mutations on top of that history afterwards is equivalent to having generated them during the simulation: it does not matter if we generate each generation’s mutations during the simulation, or afterwards. To add mutations after the fact, we need to know the genealogical trees relating all sampled individuals at each position along the genome. Combined with ancestral genotypes and the origins of new mutations, these trees completely specify the genomic sequence of any individual in the population at any time. To obtain this information, we record from forward simulation the *population pedigree* – the complete history of parent-offspring relationships of an entire population going back to a remote time – and the genetic outcomes of each ancestral meiosis, periodically discarding all information irrelevant to the genetic history of the extant population. The information in this embellished pedigree is stored as a *succinct tree sequence* (or, for brevity, “tree sequence”), which contains all the information necessary to construct the genealogical tree that relates each individual to every other at each position on the genome.

The idea of storing genealogical information to speed up simulations is not new. It was implemented in AnA-FiTS (Aberer and Stamatakis 2013), but without the critical step of discarding irrelevant genealogical information. Padhukasahasram et al. (2008) obtained impressive speedups for a Wright-Fisher simulation by keeping track of genealogies over the preceding 8 generations and only tracking neutral genotypes for those segments having descendants across this window. Our approach is similar, but uses genealogies across the entire duration of the simulation. The embellished pedigree is equivalent to the *ancestral recombination graph*, or ARG (Griffiths 1991; Griffiths and Marjoram 1997), which has been the subject of substantial study (Marjoram and Wall 2006; Wilton et al. 2015; Wiuf and Hein 1997, 1999). However, it is unclear if an ARG-based approach would share the computational advantages of the data structures we use here (Kelleher et al. 2016).

In this paper, we describe a storage method for *succinct tree sequences* (and hence, genome sequence) as well as an algorithm for simplifying these. The data structure is *succinct* in the sense that its space usage is close to optimal, while still allowing efficient retrieval of information (see, e.g., Gog et al. (2014)). We also describe how these tools can efficiently record, and later process, the embellished population pedigree from a forwards-time simulation. While providing substantial savings in computational time and space, our methods provide in principle much more information than simply simulating the genomes - the tree sequence encodes all marginal genealogies of individuals living at the end of the simulation. These marginal genealogies enable fast data storage and processing, but also provide additional information that can be used to better understand the notoriously complex dynamics of population genetics. Although we were motivated by a need for more efficient genomic simulations, these tools may prove more widely useful. This work originated as improvements to the algorithmic tools and data structures in the coalescent simulator msprime; the software tools described here for working with tree sequences are referred to as tskit; they are currently bundled with the Python package msprime, but will soon be separately available as a Python API and an embeddable C library.

## Results

The strategy described above is only of interest if it is computationally feasible. Therefore, we begin by benchmarking the performance improvement achieved by this method, implemented using the forwards-time simulation library `fwdpp` (Thornton 2014) and tree sequence tools implemented in tskit. Then, we describe the conceptual and algorithmic foundations for the method: (a) a format, implemented in the tskit Python API, for recording tree sequences efficiently in several *tables*; (b) an algorithm to record these tables during a forwards-time simulation; and (c) an algorithm to *simplify* a tree sequence, i.e., remove redundant information. Finally, we analyze the run time and space complexity of our general-purpose method.

### Simulation benchmarks

To measure the performance gains from recording the pedigree we ran simulations both with and without recording. (Although we record more than just the parent-offspring relationships of the pedigree, for brevity we refer to the method as “pedigree recording”.) All simulations used `fwdpp` to implement a discretetime Wright-Fisher population of *N* diploid individuals, simulated for 10*N* generations (details below). Simulations without pedigree recording introduced neutral mutations at a rate equal to the recombination rate, so *μ* = *r*, where *μ* and *r* are the expected per-generation number of mutations per gamete and recombination breakpoints per diploid, respectively. Simulations with pedigree recording introduced neutral mutations at the same rate retrospectively, as described below, resulting in statistically identical simulation results. We ran simulations with different values of *N* and varied the size of the genomic region according to the scaled recombination parameter *ρ* = 4*Nr*.

Deleterious mutations were introduced at rate *ρ*/100 per generation, drawing scaled selection coefficients (2*Ns*) from a Gamma distribution with a mean of −5 and a shape parameter of 1. This distribution of fitness effects results in thousands of weakly-deleterious mutations segregating in the population, many of which drift to intermediate frequencies. The case of many mutations with selection is a non-trivial violation of exchangeability assumptions of the coalescent (Neuhauser and Krone 1997). Therefore, these selected mutations must be explicitly tracked in our forward simulation and the time savings due to pedigree recording come from not having to record *neutral* mutations.

Pedigree tracking dramatically reduced runtimes, as shown in Figure 1, producing a relative speedup of up to around 50 fold relative to standard simulations that track neutral mutations (Figure 2). Pedigree tracking results in greater relative speedups for larger *N* and we observe increasing relative speedups as 4*Nr* increases for a given *N* (Figure 2). Importantly, runtimes are approximately linear in region size *ρ* when pedigree tracking (partially obscured by the log scale of the horizontal axis in Figure 1). In a more limited set of neutral simulations we found the same qualitative behavior, and a numerically larger speedup by using pedigree tracking (see Appendix A).

**Figure 1:**
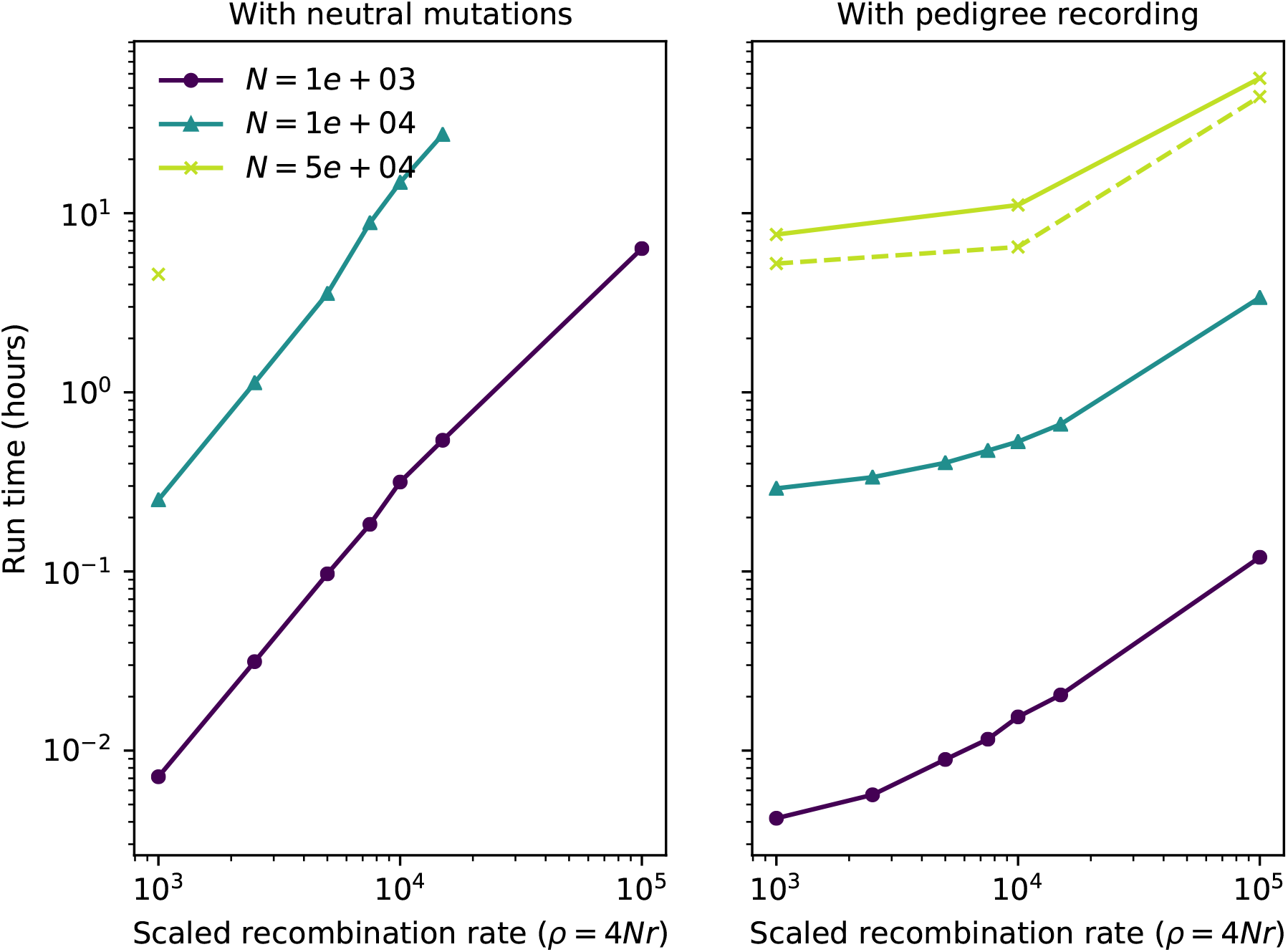
Total run time per single simulation replicate as a function of region length. Line color represents different diploid population sizes (*N*). The left figure shows run times for standard simulations including neutral mutations. The right column shows run times of simulations that recorded the pedigree and added neutral mutations afterwards. The dashed line in the right panel shows results for an implementation using `fwdpy11` where the pedigree simplification steps were handled in a separate thread of execution and fitness calculations were parallelized across four cores. Simulations with *N* = 5 × 10^4^ timed out for region sizes larger than 10^3^.

**Figure 2:**
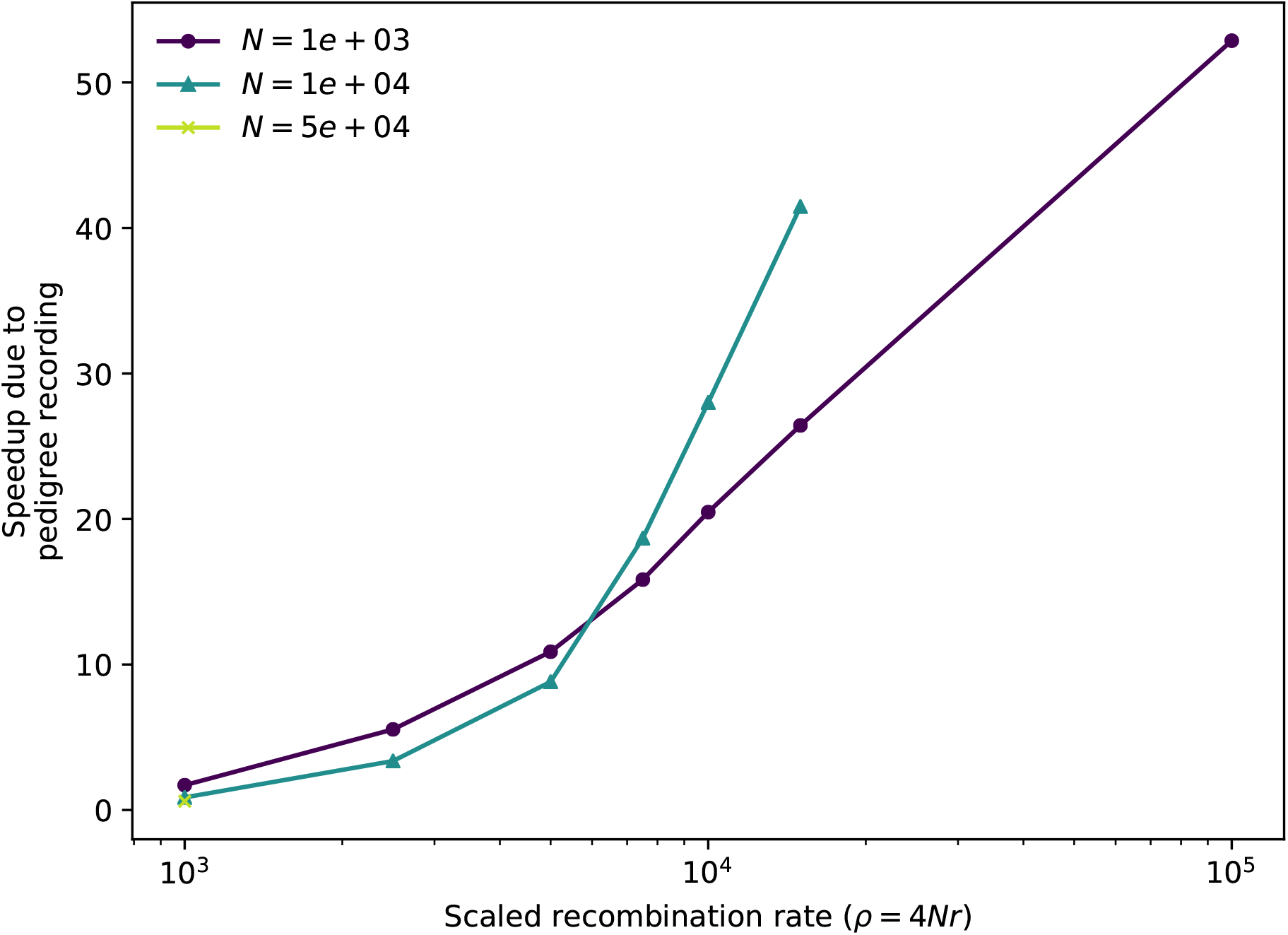
Relative speedup of simulations due to pedigree recording. Each line shows the ratio of total run times of standard simulations to those of simulations with pedigree recording. Data points are taken from Figure 1 for simulations that ran to completion in both cases.

In our implementation, simulations with pedigree recording used substantially more RAM than simple forward simulations (see Appendix B). This is unsurprising: unsimplified tree sequences grow quickly, and so storing history can use arbitrarily much memory. However, this is not a requirement of the method, only a straightforwards consequence of a speed-memory tradeoff: the amount of required memory is mostly determined by the interval between simplification steps, but less frequent simplification reduces overall computation time (see Appendix C). In fact, our method could in some situations *reduce* the amount of memory required, if memory usage in the forwards simulation was dominated by the cost of maintaining neutral genetic variants.

### Tables for succinct tree sequences

We now explain what we actually did to achieve this 50 × speedup. The “pedigree recording” simulations above recorded information about each new individual in a collection of tables that together define a *succinct tree sequence* (or, simply “tree sequence”). A tree sequence is an encoding for a sequence of correlated trees, such as those describing the history of a sexual population. Tree sequences are efficient because branches that are shared by adjacent trees are stored once, rather than repeatedly for each tree. The topology of a tree sequence is defined via its *nodes* and *edges*, while information about variants is recorded as *sites* and *mutations*; we give an example in Figure 3. This formulation is derived from the “coalescence records” encoding of tree sequences (Kelleher et al. 2016), normalised to remove redundancy and generalised to include a more general class of tree topologies.

**Figure 3:**
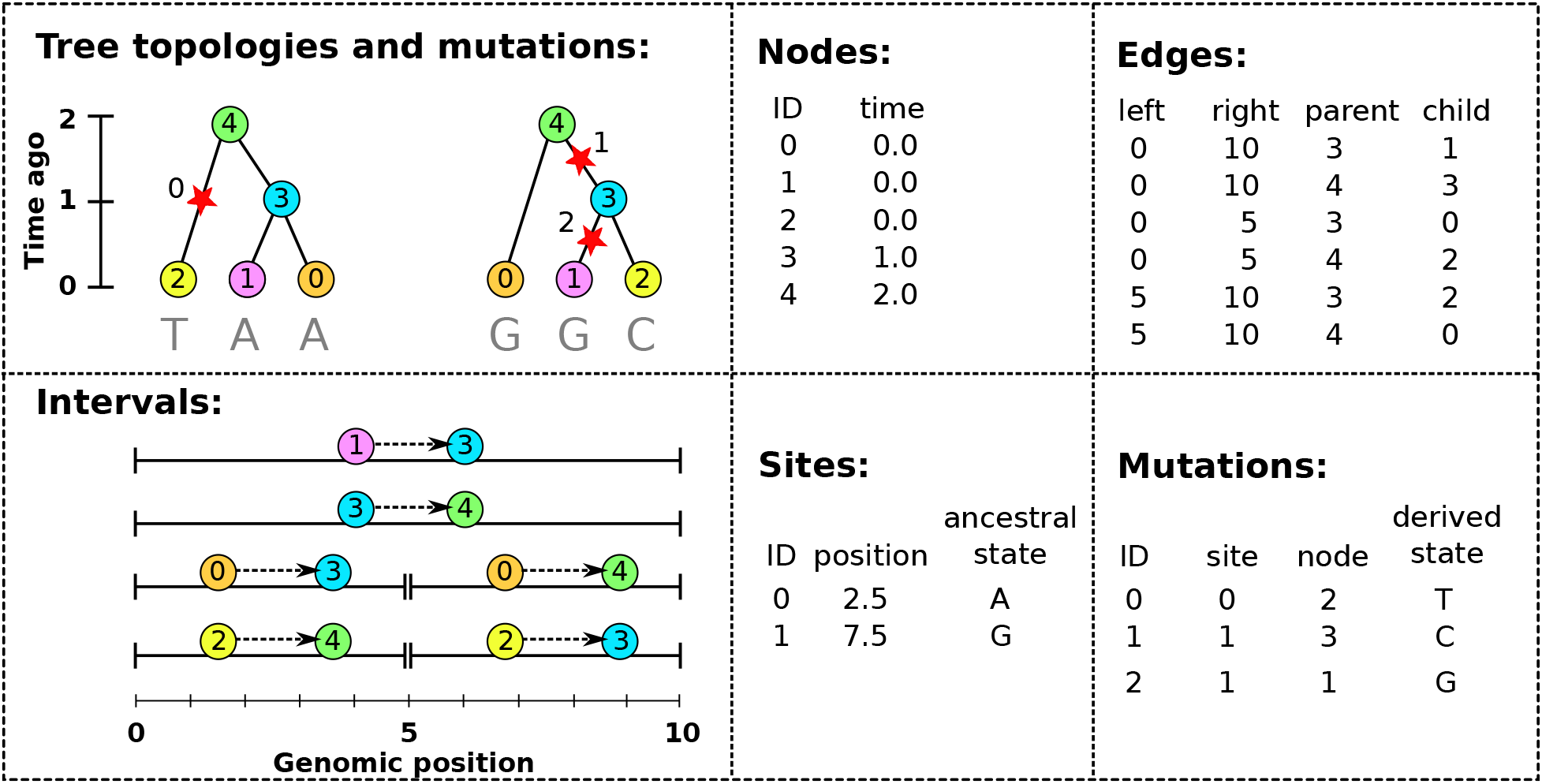
An example tree sequence with three samples over a chromosome of length 10. The leftmost panels show the tree sequence pictorially in two different ways: (top) a sequence of tree topologies; the first tree extends from genomic position 0 to 5, and the second from 5 to 10; and (bottom) the edges that define these topologies, displayed over their corresponding genomic segment (for instance, the edge from node 2 to node 4 is present only on the interval from 0 to 5). The remaining panels show the specific encoding of this tree sequence in the four tables (nodes, edges, sites and mutations).

The *nodes* of a tree sequence correspond to the vertices in the individual genealogies along the sequence. Each node refers to a specific, distinct ancestor, and so has a unique “time”, thought of as the node’s birth time, which determines the height of any vertices the node is associated with. (Note that since each node time is equal to the amount of time since the birth of the corresponding parent, time is measured in clock time, not in meioses.) The example of Figure 3 has five nodes: nodes 0, 1 and 2 occur at time 0 and are the *samples*, while nodes 3 and 4 represent those ancestors necessary to record their genealogy, who were born one and two units of time in the past, respectively.

The *edges* define how nodes relate to each other over specific genomic intervals. Each edge records the endpoints [*ℓ, r*) of the half-open genomic interval defining the spatial extent of the edge; and the identities *p* and *c* of the parent and child nodes of a single branch that occurs in all trees in this interval. The spatial extent of the edges defining the topology of Figure 3 are shown in the bottom left panel. For example, the branch joining nodes 1 to 3 appears in both trees, and so is recorded as a single edge extending over the whole chromosome. It is this method of capturing the shared structure between adjacent trees that makes the tree sequence encoding compact and algorithmically efficient.

Recovering the sequence of trees from this information is straightforward: each point along the genome at which the tree topology changes is accompanied by the end of some edges and the beginning of others. Since each edge records the genomic interval over which a given node inherits from a particular ancestor, to construct the tree at a certain point in the genome we need only retrieve all edges overlapping that point and construct the corresponding tree. To modify the tree to reflect the genealogy at a nearby location, we simply remove those edges whose intervals do not overlap that location, and add those new edges whose intervals do. Incidentally, this property that edges naturally encode *differences* between nearby trees (e.g., as “subtree prune and regraft” moves) allows for efficient algorithms to compute statistics of the genome sequence that take advantage of the highly correlated nature of nearby trees (Kelleher et al. 2016).

Given the topology defined by the nodes and edges, *sites* and *mutations* encode the sequence information for each sample in an efficient way. Each site records two things: its position on the genome and an ancestral state. For example, in Figure 3 we have two sites, one at position 2.5 with ancestral state ‘A’ and the other at position 7.5 with ancestral state ‘G’. If no mutations occur at a given site, all nodes inherit the ancestral state. Each mutation records three things: the site at which it occurs, the first node to inherit the mutation, and the derived state. Thus, all nodes below the mutation’s node in the tree will inherit this state, unless further mutations are encountered. Three mutations are shown in Figure 3, illustrated by red stars. The first site, in the left-hand tree, has a single mutation, which results in node 2 inheriting the state ‘T’. The second site, in the right hand tree, has two mutations: one occurring over node 3 changing the state to ‘C’, and a back mutation over node 1 changing the state to ‘G’.

This encoding of a sequence of trees and accompanying mutational information is very concise. To illustrate this, we used `msprime` to simulate 500,000 samples of a 200 megabase chromosome with humanlike parameters: *N_e_* = 10^4^ and per-base mutation and recombination rates of 10^−8^ per generation. This resulted in about 1 million distinct marginal trees and 1.1 million infinite-sites mutations. The HDF5 file encoding the node, edge, site and mutation tables (as described above) for this simulation consumed 157MiB of storage space. Using the tskit Python API, the time required to load this file into memory was around 1.5 seconds, and the time required to iterate over all 1 million trees was 2.7 seconds. In contrast, recording the topological information in Newick format would require around 20 TiB and storing the genotype information in VCF would require about 1 TiB (giving a compression factor of 144,000 in this instance). Working with either the Newick or VCF encoding of this dataset would likely require several days of CPU time just to read the information into memory.

#### Validity of a set of tables

Given a set of node and edge tables as described above, there are only two requirements that ensure the tables describe a valid tree sequence. These are:

1. Offspring must be born after their parents (and hence, no loops).
2. The set of intervals on which each individual is a child must be disjoint.

A pair of node and edge tables that satisfy these two requirements is guaranteed to uniquely describe at each point on the genome a collection of directed, acyclic graphs – in other words, a forest of trees. For some applications it is necessary to check that at every point there is only a *single* tree. Checking this is more difficult, but is implemented in `tskit`’s API. For efficiency, `tskit` makes several other sortedness requirements on the tables, that are not necessarily satisfied by tables emitted by a forwards-time simulation. `tskit`’s API includes tools to rectify this by first sorting and then using the `simplify` algorithm described below, which works on sorted tables and is guaranteed to produce a valid, `tskit`-ready tree sequence.

### The tskit Tables API

The facilities for working with succinct tree sequences are implemented as part of the `tskit` Python API, which provides a powerful platform for processing tree topology and mutation data. The portions of `tskit` that we discuss here are dedicated to tree sequence input and output using simple tables of data, as described above, so we refer to this as the “Tables API”.

The Tables API is primarily designed to facilitate efficient interchange of data between programs or between different modules of the same program. We adopted a ‘columnar’ design, where all the values for a particular column are stored in adjacent memory locations. There are many advantages to columnar storage – for example, since adjacent values in memory are from the same column, they tend to compress well, and suitable encodings can be chosen on a per-column basis (Abadi et al. 2006). A particular advantage of this approach is that it enables very efficient copying of data, and in principle zero-copy data access (where a data consumer reads directly from the memory of a producer). Our implementation efficiently copies data from Python as a NumPy array (Walt et al. 2011) into the low-level C library used to manipulate tree sequences. This architecture allows for data transfer rates of gigabytes per second (impossible under any text-based approach), while retaining excellent portability. NumPy’s array interface provides a great deal of flexibility and efficiency, and makes it straightforward to transfer data from sources such as HDF5 (The HDF Group 1997-2018) or Dask (Dask Development Team 2016). For small scale data and debugging purposes, a simple text based format is also supported.

The tskit Python Tables API provides a general purpose toolkit for importing and processing succinct tree sequences, and a collection of tutorials are being developed at https://github.com/tskit-dev/tutorials. Interoperation with Python simulators is then straightforward. The implementation we benchmark here uses `pybind11` (https://github.com/pybind/pybind11/) to interface with the `fwdpp C++` API (Thornton 2014). No modifications were required to the `fwdpp` code base; rather, we simply need to bookkeep parent/offspring labels, and perform simple processing of the recombination breakpoints from each mating event to generate node and edge data. This information is then periodically copied to the tskit Tables API, where it is sorted and simplified.

#### Flexibility

To demonstrate the flexibility provided by the Tables API and provide an implementation that decouples forward simulation internals from transfer of data to `tskit`, we also implemented a version of the simulations described in “Simulation benchmarks” separately in Python, described in Appendix D. In this proof-of-concept implementation, the simulation engine (we use `simuPOP`, Peng and Kimmel (2005)) invokes callbacks at critical points of the simulation, and we infer nodes and edges from the information that is provided. Rows are appended to the tables one-by-one, and the tables are periodically sorted and simplified to control memory usage. Benchmarking results from this implementation are shown (alongside results from `fwdpp`) for simulations without selection in Appendix A: a relatively modest speedup of around 5× is achieved, likely due to increased overhead.

### Recording the pedigree in forwards time

To record the genealogical history of a forwards time simulation, we need to record two things for each new chromosome: the birth time; and the endpoints and parental IDs of each distinctly inherited segment. These are naturally stored as the *nodes* and *edges* of a tree sequence. To demonstrate the idea, we write out in pseudocode how to run a neutral Wright-Fisher simulation that records genealogical history in this way. The simulation will run for *T* generations, and has *N* haploid individuals, each carrying a single chromosome of length L. For simplicity, we sample exactly one crossover per generation. Note that the *table recording* portion of the algorithm does not depend on the Wright-Fisher nature of the population simulation; next we will describe how to record tables from *any* simulation.

We use 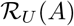 to denote an element of the set A chosen uniformly at random (and all such instances are independent). Given a node table 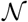, the function 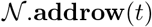 adds a new node to the table 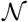 with time *t* and returns the ID of this new node. Similarly, the function ℰ.**addrow**(*ℓ,r,p,c*) adds a new edge (feft, right, parent, child) to the edge table ℰ. The function **simplify** 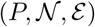 (described below) simplifies the history stored in the tables 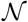 and *ℰ* to the minimal information required to represent the genealogies of the list of node IDs *P*; after simplification the nodes appearing in *P* are relabeled (0, 1, …, |*P*| − 1). A step-by-step explanation follows the pseudocode.

#### Algorithm W.

(*Forwards-time tree sequence*). Simulates a randomly mating population of *N* haploid individuals with chromosome of length *L* for *T* generations, and returns the node and edge tables (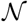 and ℰ) recording the simulated history. In each generation, the current population is stored in *P*, while produced offspring are placed in *P*′. The tables are simplified every s generations, removing genealogical information from 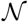 and ℰ irrelevant to the current population.

**W1**. [Initialisation.] Set 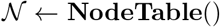, ℰ ← **EdgeTable**(), *t* ← *T*, and *j* ← 0. For 0 ≤ *k* < *N*, set 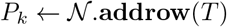.
**W2**. [Generation loop head: new node.] Set 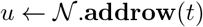 and 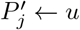.
**W3**. [Choose parents.] Set 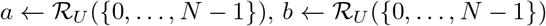 and 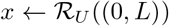.
**W4**. [Record edges.] Call ℰ.**addrow**(0, *x, P_a_, u*) and ℰ.**addrow**(*x, L, P_b_, u*).
**W5**. [Individual loop.] Set *j* ← *j* + 1. If *j* < *N* go to W2. Otherwise, if *t* mod *s* ≠ 0 go to W7.
**W6**. [Simplify.] Call **simplify**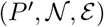, and set 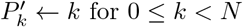 for 0 ≤ *k* < *N*.
**W7**. [Generation loop.] Set *t* ← *t* − 1. If *t* = 0 terminate. Set *P* ← *P*′, *j* ← 0, and go to W2.

We begin in W1 by creating new node and edge tables, and setting our population *P* (a vector of *N* node IDs) to the initial population. This initial population is a set of *N* nodes with birth time *T* generations ago. We also initialise our generation clock *t* and individual index *j*. Step W2 replaces the *j*^th^ individual (with node ID *P_j_*) by creating a new node with birth time *t* (and ID *u*). In step W3 we determine the new node’s ancestry by choosing two indexes *a* and *b* uniformly, giving us parental IDs *P_a_* and *P_b_*, and choose a chromosomal breakpoint *x*. We record the effects of this event by storing two new edges: one recording that the parent of node *u* from 0 to *x* is *P_a_*, and another recording that the parent of *u* from *x* to *L* is *P_b_*. Step W5 then iterates these steps for each of the N individuals for each generation. At the end of a generation, we then check if we need to simplify (done every s generations). If simplification is required, we do this in step W6 by calling the simplify function on the node and edge tables with the current set of population IDs *P*′ as the samples. This updates the tables in-place to remove all redundant information, and remaps the specified sample IDs to 0, …, *N* − 1 in the updated tables. Hence, we set our current population IDs to 0, … *N* − 1 after simplify has completed. Step W7 loops these steps until the required number of generations have been simulated.

This algorithm records only topological information about the simulated genealogies, but it is straight-forward to add mutational information. Mutations that occur during the simulation can be recorded by simply storing the node in which they first occur, the derived state, and (if not already present) the genomic position of the site at which it occurs. This allows selected mutations, that the forwards time simulation must generate, to be recorded in the tree sequence. Neutral mutations can be generated after the simulation has completed, thus avoiding the cost of generating the many mutations that are lost in the population. This is straightforward to do because we have access to the marginal genealogies.

Figure 4 shows an example of a marginal genealogy produced by a forwards-time Wright-Fisher process like Algorithm W. On the left is the tree showing all the edges output by the simulation, while on the right is the minimal tree representing the ancestry of the current set of samples. Clearly there is a great deal of redundancy in the topological information output by the simulation. This redundancy comes from two sources. First, there are a large number of nodes in the tree that have only one child. In Algorithm W we do not attempt to identify coalescence events, but simply record all parent-child relationships in the history of the population. As such, many of these edges will record the simple passing of genealogical information from parent to child and only some small subset will correspond to coalescences within the marginal trees. The second source of redundancy in the (unsimplified) output of Algorithm W is due to the fact that lineages die out: a large number of individuals in the simulation leave no descendants in the present day population. Node 26 in Figure 4a, for example, leaves no ancestors in the current population, and so the entire path tracing back to its common ancestor with 27 is redundant.

**Figure 4:**
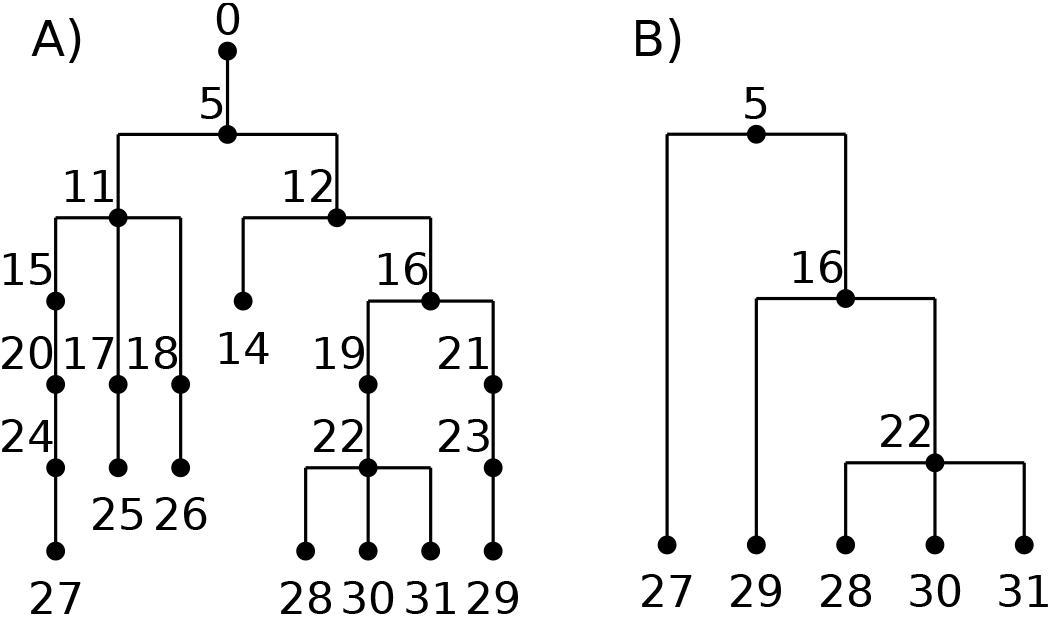
An example of a marginal genealogy from a Wright-Fisher simulation with *N* = 5. (**A**) the original tree including all intermediate nodes and dead-ends, and (**B**) the minimal tree relating all of the currently-alive individuals (27–31).

#### Essential steps in tree sequence recording

Since a tree sequence can record the history of genetic inheritance in any situation (requiring only unambiguous inheritance and no time travel), any individual-based population genetics simulator can maintain a tree sequence with only a little bookkeeping. We have furthermore provided several tools to minimize this bookkeeping, only requiring one-way output at the birth of each new individual. Concretely, to record a tree sequence, including mutations, a simulator must record for each new genome (so, twice for each new diploid individual):

- the birth time of the genome in the Node Table,
- the segments the genome inherits from its parental genomes in the Edge Table,
- the locations of any new mutations in the Site Table,
- and the derived state of these new mutations in the Mutation Table (as well as the identity of this genome the mutations appeared in).

Each of these can be simply appended to the ends of the respective tables. Besides this, simplification should be run every once in a while (e.g., every 100 generations). Before simplification, time in the Node Table must be translated to “time ago” if it is not already. (This was avoided in Algorithm W since there t denoted “time until the end of the simulation”.) There are also several “cleaning” steps, for which we provide functions in the Tables API: *sorting* according to several criteria for algorithmic efficiency; and removing any duplicate sites from the Site Table. After simplification, since **simplify**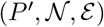 results in the *N* individuals in *P*′ being relabeled in tskit’s tables as 0, 1, …, *N* − 1, there must be bookkeeping that keeps in sync the individual IDs as recorded by the simulator with the node IDs recorded in the tables.

In other words, to record a tree sequence, a simulation needs only to know (a) which genomes recombined to produce each new genome, and how; (b) the locations and results of any new mutations on each genome; and (c) the identities of every currently alive individual at each time simplification occurs.

#### Storing metadata

Applications may also want to store more information not fitting into an existing column of the tables, such as the selection coefficient of a mutation, or the sex of an individual. This (and, indeed, arbitrary information) can be stored in the metadata columns present in Node, Site, and Mutation tables.

### Tree sequence simplification

It is desirable for many reasons to remove redundant information from a tree sequence. To formalize this: suppose that we are only interested in a subset of the nodes of a tree sequence (which we refer to as our ‘samples’), and wish to reduce this input tree sequence to the smallest one that still completely describes the history of the specified samples, having the following properties:

1. All marginal trees must match the subtree of the corresponding tree in the input tree sequence that is induced by the samples.
2. Within the marginal trees, all non-sample vertices must have at least two children (i.e., unary tree vertices are removed).
3. Any nodes and edges not ancestral to any of the sampled nodes are removed.
4. There are no adjacent redundant edges, i.e., pairs of edges (*ℓ, x, p, c*) and (*x,r,p,c*) which can be represented with a single edge (*ℓ, r, p, c*).

Simplification is essential not only for keeping the information recorded by forwards simulation manageable, but also is useful for extracting subsets of a tree sequence representing a very large dataset.

We implement simplification by starting at the end of the simulation, and moving back up through history, recording in the new tree sequence only that information necessary to construct the tree sequence of the specified individuals. This process of tracing ancestry back through time in a pedigree was the motivation for Hudson’s coalescent simulation algorithm (Hudson 1983), so it is unsurprising that simplification uses many of the same tools as the implementation of Hudson’s algorithm in msprime (Kelleher et al. 2016). The main difference is that events in a coalescent simulation are random, while in our simplification algorithm they are predetermined by history. An implementation in pseudocode is provided in Appendix E, and a python implementation as supplementary information.

Conceptually, this works by (a) beginning by painting the chromosome in each sample a distinct color; (b) moving back through history, copying the colors of each chromosome to the portions of its parental chromosomes from which it was inherited; (c) each time we would paint two colors in the same spot (a coalescence), record that information as an edge and instead paint a brand-new color; and (d) once all colors have coalesced on a given segment, stop propagating it. This “paint pot” description misses some details – for instance, we must ensure that all coalescing segments in a given individual are assigned the *same* new color – but is reasonably close. Figure 5 shows an example tree sequence, before and after simplification, and Figure 6 depicts the “paint pot” state of the algorithm during the process of simplifying this tree sequence. Since the method begins with the samples and moves back through time, in the output tree sequence, the n samples will be numbered 0, 1, …, *n* − 1 and subsequent nodes will be ordered by time since birth. This is seen in the red labels of Figure 5.

**Figure 5:**
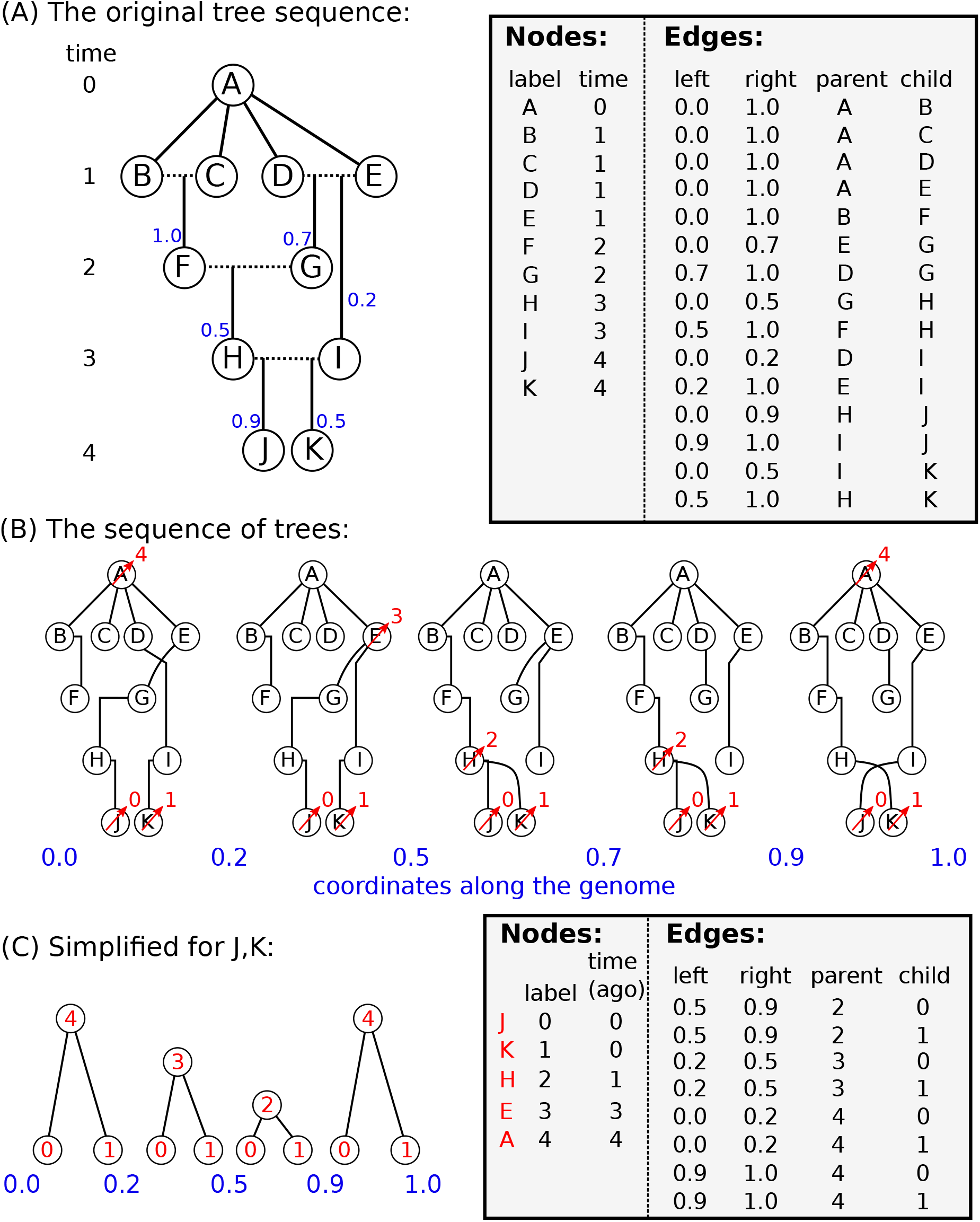
An example of tree sequence simplification. (A) The augmented pedigree diagram on the left relates the ten homologous chromosomes of five diploid individuals (BC, DE, FG, HI, and JK) to each other and to a common ancestral chromosome (**A**); dotted lines connect the two chromosomes of each individual, and solid lines lead to the products of their meioses. The corresponding tables (right) have 11 node records (one for each chromosome) and 15 edge records (one for each distinctly inherited segment). Blue numbers denote crossing over locations – for instance, *D* and *E* were parents to *G*, who inherited the left 70% of the chromosome from *E* and the remainder from *D. B, C, D*, and *E* inherit clonally from *A*. (**B**) The five distinct trees found across the chromosome (blue numbers denote locations on the chromosome). Labels after simplification are shown in red (see text). (**C**) Tables recording the tree sequence after simplification with nodes *J* and *K* as samples. The mapping from labels in the forwards time simulation to nodes in the tree sequence is shown in red.

**Figure 6:**
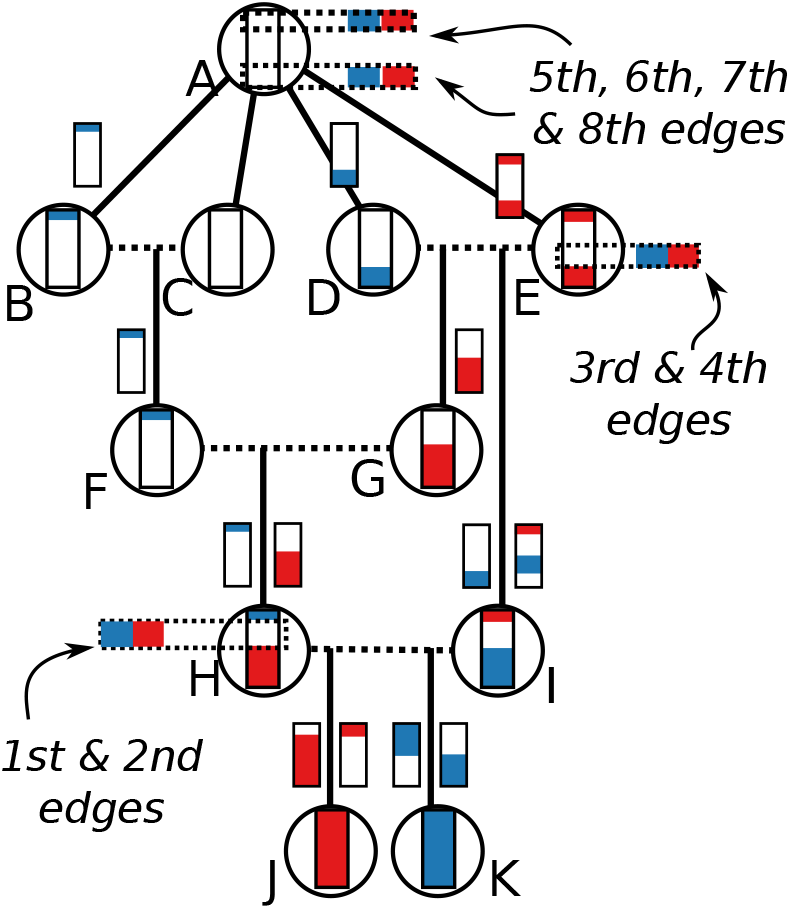
A depiction of each state of the simplification algorithm as it moves up through the embellished pedigree in the example of Figure 5A. Following the “paint pot” description in the text, we begin by coloring J and K’s genomes in red and blue respectively, then trace how these colors were inherited back up through the pedigree until they coalesce. To aid in this, the smaller colored chromosomes on either side of each solid arrow show the bits inherited from each of the two parental chromosomes, with genomic position 0.0 on the bottom and 1.0 at the top. Each time a red and a blue segment overlap, a coalescence occurs, two edges are output, and we stop propagating that segment. For instance, both J and K inherit from H between 0.5 and 0.9, which resulted in the first two edges of the simplified table of Figure 5C. Later, both inherit from E between 0.2 and 0.5, along the paths J-H-G-E and K-I-E respectively, resulting in the next two edges.

More concretely, the algorithm works by moving back through time, processing each parent in the input tree sequence in chronological order. The main state of the algorithm at each point in time is a set of ancestral lineages, and each lineage is a linked list of ancestral segments. An ancestral segment (*ℓ, r, u*) is found in a lineage if the output node u inherits the genomic interval [*ℓ, r*) from that lineage (and so u corresponds to a “color” in the description above). We also maintain a map from input nodes to lineages. Crucially, the time required to run the algorithm is linear in the number of edges of the input tree sequence.

### Sequential simplification and prior history

Any simulation scheme that records data into tables, as Algorithm W does, has its genealogical history available at any time as a tree sequence. This has two additional advantages: First, simplification can be run periodically through the simulation, if we take the set of samples to be the entire currently alive population. This is important in practice as it keeps memory usage from growing linearly (and quickly) with time. Second, the simulation can be *begun* with a tree sequence produced by some other method – for instance, by a coalescent simulation with `msprime`, providing an easy, efficient way to specify prior history. A natural question is now: how often should simplification occur? Limited testing (described in Appendix C) found that different simplification intervals affect run times by approximately 25%, with the lowest run time occurring when simplifying every 10^3^ generations. Thus, there is a memory-versus-speed tradeoff – simplifying more often would keep fewer extinct nodes and edges in memory.

### Computational complexity

Figures 1 and 2 show that this method can dramatically improve simulation performance in practice - but, how does it perform in theory? Both computational time and storage space are depend mostly on the number of *mutations* and *edges* in the tree sequence. The key quantity to understand for this will be the total “area” of the tree sequence, which is the sum of the lengths of all ancestral genomic segments that some, but not all, of the present population has inherited. It can be found by summing the product of segment length (left minus right coordinates) and edge length (difference in birth times between parent and child), across all edges. This area is also equal to the sum of the total lengths of all marginal trees (i.e., the trees describing inheritance at each position on the genome), so can be computed as the sequence length multiplied by the mean marginal tree length. Since we analyze tree sequences arising from a Wright–Fisher model, statistical properties of a marginal tree are fairly well-described by coalescent theory. Similar arguments to those below go back at least to Watterson (1975), who also explicitly computed smaller order corrections relevant to whole-population genealogies of the Wright–Fisher model. The arguments below are mostly self-contained, but for an introduction to coalescent theory, including the basic facts used below, see Wakeley (2005).

First: how much memory do simplified tree sequences require? Consider a simulation of a Wright-Fisher population of *N* haploid individuals using Algorithm W for *T* generations. Since every chromosome inherits material from both parents, without simplification this would produce tables of *NT* nodes and 2*NT* edges. After simplification, we are left with the tree sequence describing the history of only the current generation of *N* individuals, back to either *T* generations ago, or their common ancestor, whichever comes first. The tree sequence must store edges describing the leftmost marginal tree, which requires at most 2*N* − 2 edges. Then, each time the marginal tree changes along the sequence, four edges end and four new edges begin (except for changes affecting the root, which require fewer; see Kelleher et al. (2016)). Suppose that 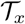 is the marginal tree at genomic position *x*, and write 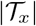 for the total length of the tree. For the tree at nearby position *x* + *dx* to be different, there must have been a crossing-over between *x* and *x* + *dx* in one of the 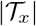 meioses that gave birth to those individuals. (Recall that these “individuals” are haploid.) Since we measure distance along the genome so that length is equal to the expected number of crossing-overs per generation, the expected distance until the next crossing-over is 1/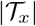. If every crossing-over changed the marginal tree, then this would imply, for consistency, that the expected total number of times that the marginal tree changes along the genome is equal to the total area of the tree sequence (as defined above). Not every such crossing-over changes the marginal tree, but most do, so the total area of the tree sequence, multiplied by four, gives an upper bound on the expected number of edges in the tree sequence beyond those describing the leftmost tree. Since we are considering a chromosome of length 1, the expected total area is equal to the mean marginal tree length, as above.

If *T* is large relative to *N*, so that all marginal trees have a single root with high probability, then coalescent theory tells us that the expected total length of the branches of a marginal tree back to the most recent common ancestor is approximately 2*N* log(*N*). Therefore, the tree sequence describing the entire population is expected to need no more than 2*N* + 8*N*log(*N*) edges. Not every new edge derives from a never-before-seen node, but the number of nodes is at most equal to the number of edges plus the sample size. Therefore, we would need *O*(*N*^2^) space to store the complete history of the simulation, but only *O*(*N*log *N*) to store the history that is relevant to the extant population.

What if *T* is smaller: how many of the resulting 2*NT* edges are required after simplification? In other words, how fast does the information in the pedigree become irrelevant? Now, we need to compute the expected total length of all branches in a coalescent tree up until time *T* (or the common ancestor, whichever comes first). Again, coalescent theory tells us that the expected length of time for which a coalescent tree has k lineages is 2*N*/(*k*(*k* − 1)) = 2*N*(1/(*k* − 1) − 1/*k*) generations. Since the tree has *k* branches over this period, it is expected to contribute 2*N*/(*k* − 1) to the total tree length. By summing over *n* < *k* ≤ *N*, the *N* tips of a tree are expected to descend from only *n* lineages around 2(*N*/*n* − 1) generations ago. Inverting this relationship between time and number of roots implies that a marginal tree cut *T* units of time ago is expected to have around *r*(*T*) roots, where *r*(*T*) = 2*N*/(*T* + 2). The total tree length over this time is 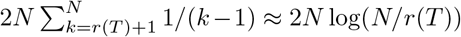, since 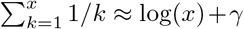. This is a crude estimate for several reasons: first, we should not count the branch leading to the root of the tree (i.e., when *r*(*T*) = 1), and second, this does not account for the discrete nature of the Wright-Fisher model. Nonetheless, this leads as above to an upper bound on the number of edges of

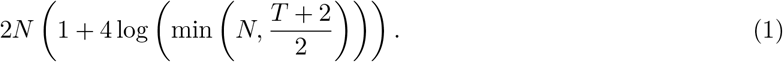

This implies that the number of edges required to store the last *T* generations of history for the entire chromosome of a population of size *N* grows as *O*(*N* log *T*) – proportionally to N at first but rapidly tapering off. The bound holds up reasonably well in practice, as shown in Figure 7.

**Figure 7:**
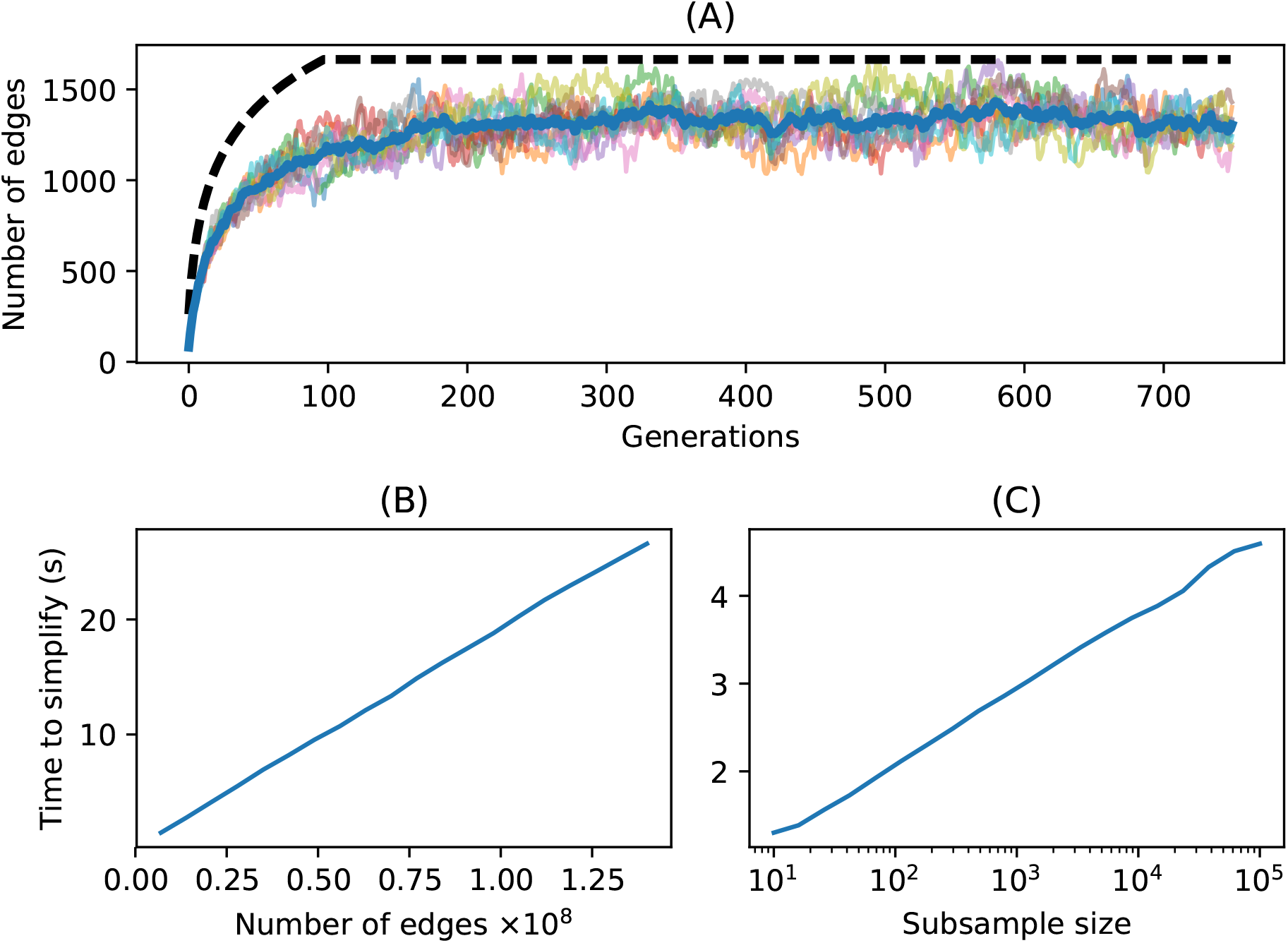
Time and space complexity of simplify. (A) Number of edges in the simplified tree sequence for 10 replicate Wright–Fisher simulations with *N* = 100 as a function of number of generations. Each line is one simulation, the heavy blue line gives the average, and the dashed line is the upper bound of equation (1). (B) Time required to simplify the first *k* edges of a large (4.2GiB) unsimplified tree sequence produced by a forwards-time simulation plotted against *k*. The time scales linearly with the number of input edges. (C) Time required to simplify the tree sequence resulting from a coalescent simulation of 500,000 samples of a 200 megabase human chromosome to a random subsample of *n* samples, plotted against *n* (note the log scale; the time scales logarithmically with *n*).

What about mutations? Forwards-time generation of infinite-sites mutations with total mutation rate per generation *μ* would produce around *μNT* mutations (and the same number of sites), simply because there was a total of NT meioses. Since mutations that are retained after simplification are precisely those that fall on the marginal trees, the number of mutations is proportional to the total area of the tree sequence. By the same argument as for the number of edges, this will result in around *μ*2*N* log(*N*) mutations. With *N* = 2× 10^4^ and *T* = 10*N*, this implies that adding neutral mutations to the simplified tree sequence reduces the number of mutations that must be generated by a factor of 10,000. This could result in substantial time savings, even without considering the computational burden of propagating mutations forwards across generations.

Since each mutation is stored only as a single row in the mutation table, and at most one row in the site table, the space required for *M* mutations is *O*(*M*); combined with the *O*(*N*log*N*) storage for edges and nodes of a simplified tree sequence, this implies that the full results of a simulation of *N* individuals having M mutations can be stored in *O*(*N* log *N*+*M*) space. In simulations of whole chromosomes, there are typically a small, bounded number of mutations per chromosome per generation, so *M* will be *O*(*N*log *N*) as well.

How does the computation *time* required for simplification scale? Simply because it must process each edge, the simplification algorithm is at least linear in the number of edges of the input tree sequence. Empirically the algorithm is exactly linear, as seen in Figure 7B, which shows the time required to simplify increasingly large subsets of a large tree sequence. When simplifying the result of a forwards-time sequence, the number of edges is the main contributing factor. Suppose on the other hand we want to simplify an already-minimal but large tree sequence with *N* nodes to a subsample of size *n*. How does the required time scale with *n*? In this case, the computation is dominated by the size of the output tree sequence, which grows with log(*n*), as shown in Figure 7C.

## Discussion

In this paper, we have shown that storing pedigrees and associated recombination events in a forwards-time simulation not only results in having available a great deal more information about the simulated population, but also can speed up the simulation by orders of magnitude. To make this feasible, we have described how to efficiently store this information in numerical tables, and have described a fundamental algorithm for simplification of tree sequences. Conceptually, recording of genealogical and recombination events can happen independently of the details of simulation; for this reason, we provide a well-defined and well-tested API in Python for use in other code bases (a C API is also planned).

The tree sequences produced by default by this method are very compact, storing genotype *and* genealogical information in a small fraction of the space taken by a compressed VCF file. The format also allows highly efficient processing for downstream analysis. Efficient processing is possible because many statistics of interest for population genetics are naturally expressed in terms of tree topologies, and so can be quickly computed from the trees underlying the tree sequence format. For example, pairwise nucleotide diversity *π*, is the average density of differences between sequences in the sample. To compute this directly from sequence data at m sites in n samples requires computing allele frequencies, taking *O*(*nm*) operations. By using the locations of the mutations on the marginal trees, and the fact that these are correlated, sequential tree algorithms similar to those in (Kelleher et al. 2016) can do this in roughly *O*(*n* + *m* +*t* log *n*) operations, where t is the number of distinct trees. The `tskit` API provides a method to compute π among arbitrary subsets of the samples in a tree sequence, which took about 0.7 seconds when applied to an example simulation of 100 megabases of human-like sequence for 200,000 samples (about 500K sites). The corresponding numeric genotype matrix required about 95GiB of RAM, and calculating *π* took about 66 seconds with NumPy.

Another attractive feature of this set of tools is that it makes it easy to incorporate *prior history*, simply by seeding the simulation with a (relatively inexpensive) coalescent simulation. This allows for incorporation of deep-time history beyond the reach of individual-based simulations. This may not even negatively affect realism, since geographic structure from times longer ago than the mixing time of migration across the range has limited effect on modern genealogies (Wilkins 2004), other than possibly changing effective population size (Barton et al. 2002; Cox and Durrett 2002).

### Other applications

The methods described here for efficiently storing tree sequences may prove useful in other fields. We have focused on the interpretation of tree sequences as the outcome of the process of recombination, but in principle, we can efficiently encode any sequence of trees which differ by subtree-prune-and-regraft operations. Since each such operation requires a constant amount of space to encode, the total space required is *O*(*n* + *t*) for *t* trees with *n* leaves (Kelleher et al. 2016). For instance, the large numbers of large, correlated trees produced by MCMC samplers used in Bayesian phylogenetics (e.g., Drummond et al. 2012) might be compactly stored as a tree sequence, which would then allow highly efficient computation of properties of the posterior distribution.

In this article, we applied our methods for storing trees to the problem of pedigree recording in a forward-time simulation. However, the method applies to any simulation scheme generating nodes and edges. For example, one could use the methods described here to generate succinct tree sequences under coalescent processes not currently implemented in `msprime`, such as the coalescent with gene conversion (Wiuf and Hein 2000), using the structured coalescent to model various forms of natural selection (Braverman et al. 1995; Kaplan et al. 1988, 1989), or the coalescent within a known pedigree. For such models, one could in principle generate tables of nodes and edges to be simplified in `tskit`. The resulting succinct tree sequence object would be in the same format as those generated by `msprime`’s simulate function, and therefore compatible with existing methods for downstream analyses.

Another application of our methods would be the case of simulating coalescent histories conditional on known pedigrees. The standard description of the Wright-Fisher coalescent averages over pedigrees. However, conditional on a realized pedigree, the distribution of coalescent times in the recent past differs from that of the unconditional coalescent (Wakeley et al. 2012). For populations with known pedigrees (*e.g*. Aguillon et al. 2017), it may be of use to simulate transmission along such pedigrees for the purpose of inference.

**A final note**: in preparing this manuscript, we debated a number of possible terms for the embellished pedigree, i.e., the “pedigree with ancestral recombination information”, the object through which each tree of a tree sequence is threaded. Etymological consensus (Liberman 2014) has “pedigree” derived from the french “pied de grue” for the foot of a crane (whose branching pattern resembles the bifurcation of a single parent-offspring relationship). An analogous term for the embellished pedigree might then be *nedigree*, from “nid de grue”, as the nest of a crane is a large jumble of (forking) branches. We thought it would be confusing to use this term throughout the manuscript, but perhaps it will prove useful elsewhere.

## Methods

We implemented simulations and the connection to tskit in `C++`, using `fwdpp` library functions and interface code using a continuum-sites model for both mutation and recombination. Simulations were run using `fwdpy11` (version 0.13.a0), a Python package based on `fwdpp` (version 0.5.7). The majority of results are presented based on a single-threaded implementation. However, we also implemented a parallelized version using Python’s `queue.Queue` to run the simplification step in a separate Python thread. Our implementation allows a maximum of four simplification intervals to be in the queue at once. This parallelized version also performed fitness calculation in parallel using two threads of execution in C++.

Code for all simulations and figures is available at https://github.com/petrelharp/ftprime_ms. These made use of the GNU Scientific Library (version 1.16 Galassi et al 2018), pybind11 (version 2.2.1 Jakob et al. 2016), and GCC (version 4.8.5). We ran all benchmarks on an Ubuntu Linux (version 16.04) system with two 2.6 GHz Intel E5-2650 CPU with hyperthreading enabled. We ran one simulation at a time and the machine was under minimal load otherwise. We used GNU parallel (Tange 2011) to kill any simulation that did not finish within 72 hours, and the Linux time command to record run time and peak memory usage of each replicate.

## Acknowledgements

Thanks to Gil McVean, Jared Galloway, Brad Shaffer, and Evan McCartney-Melstad for useful discussions. Work on this project was supported by funding from the Sloan Foundation and the NSF (under DBI-1262645) to PR; the Wellcome Trust (grant 100956/Z/13/Z) to Gil McVean; the NIH (R01GM115564) to KRT; and the USF&WS to H. Bradley Shaffer.

## A Timing and memory use with no selection

We also performed a more limited set of simulations without natural selection. For these we used `C++` as in the main text, but also provide results from our proof-of-concept implementation that uses `simuPOP` (see Appendix D).

The total run times are shown in Figure D1 and for `C++` show the same qualitative behavior as simulations with selection (Figure 1). The relative improvement due to pedigree tracking is again substantial in these simulations (Figure D2). In fact, we see more of a benefit to pedigree tracking here with `fwdpy11` than we did with selection (Figure 2). The reason is that the fitness function exits close to instantly in simulations without selection that are based on `fwdpp`, meaning that `fwdpy11` is doing little more than generating random numbers and book-keeping in Figures D1 and D2. For `simuPOP`, the results show a qualitatively similar pattern but (1) the simulations for *N* = 10000 or greater did no finish in the 72 hour limit or exceed memory limits (more than 500 GB), and (2) the magnitude of the speedup is much less. The total RAM use for simulations without selection is shown in Figure D3.

**Figure D1:**
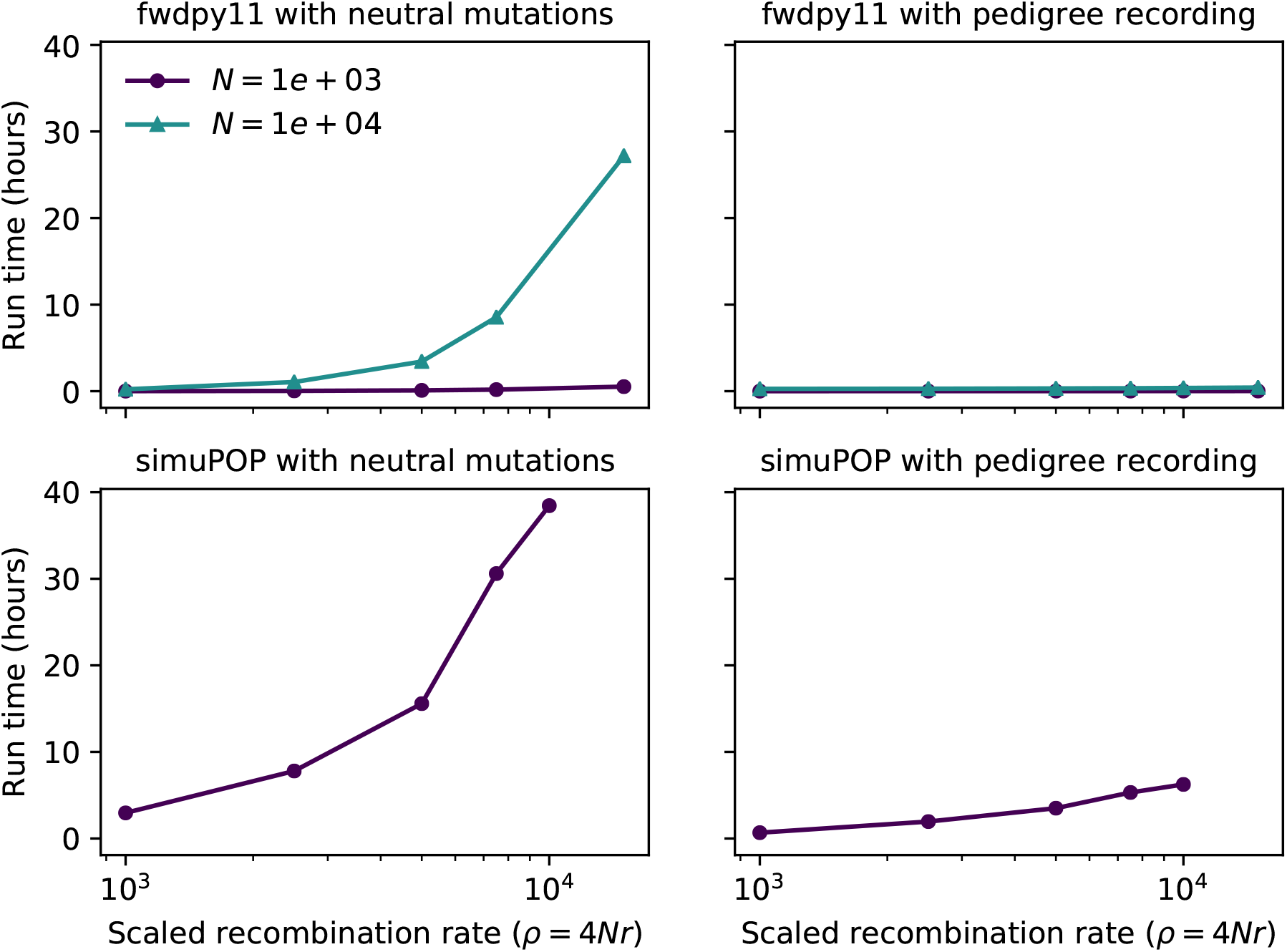
Total run time for a single simulation replicate as a function of region length, measured as the scaled recombination parameter *ρ* = 4*Nr*. The simulations here involve no natural selection.

**Figure D2:**
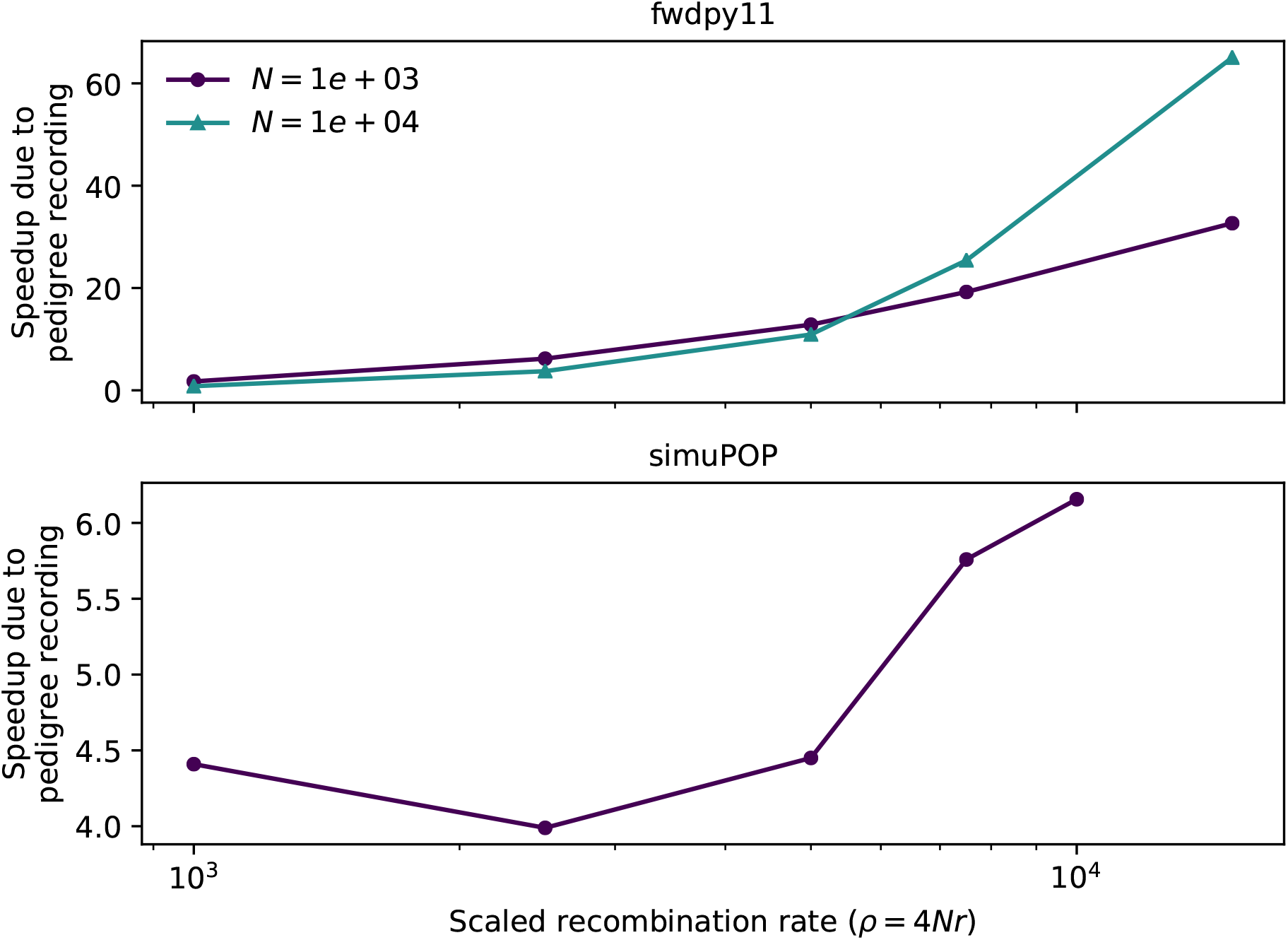
Relative performance improvement due to pedigree tracking for simulations without selection. Data points are taken from Figure D1 and show the ratio of run times tracking neutral variation over the run times tracking the pedigree.

**Figure D3:**
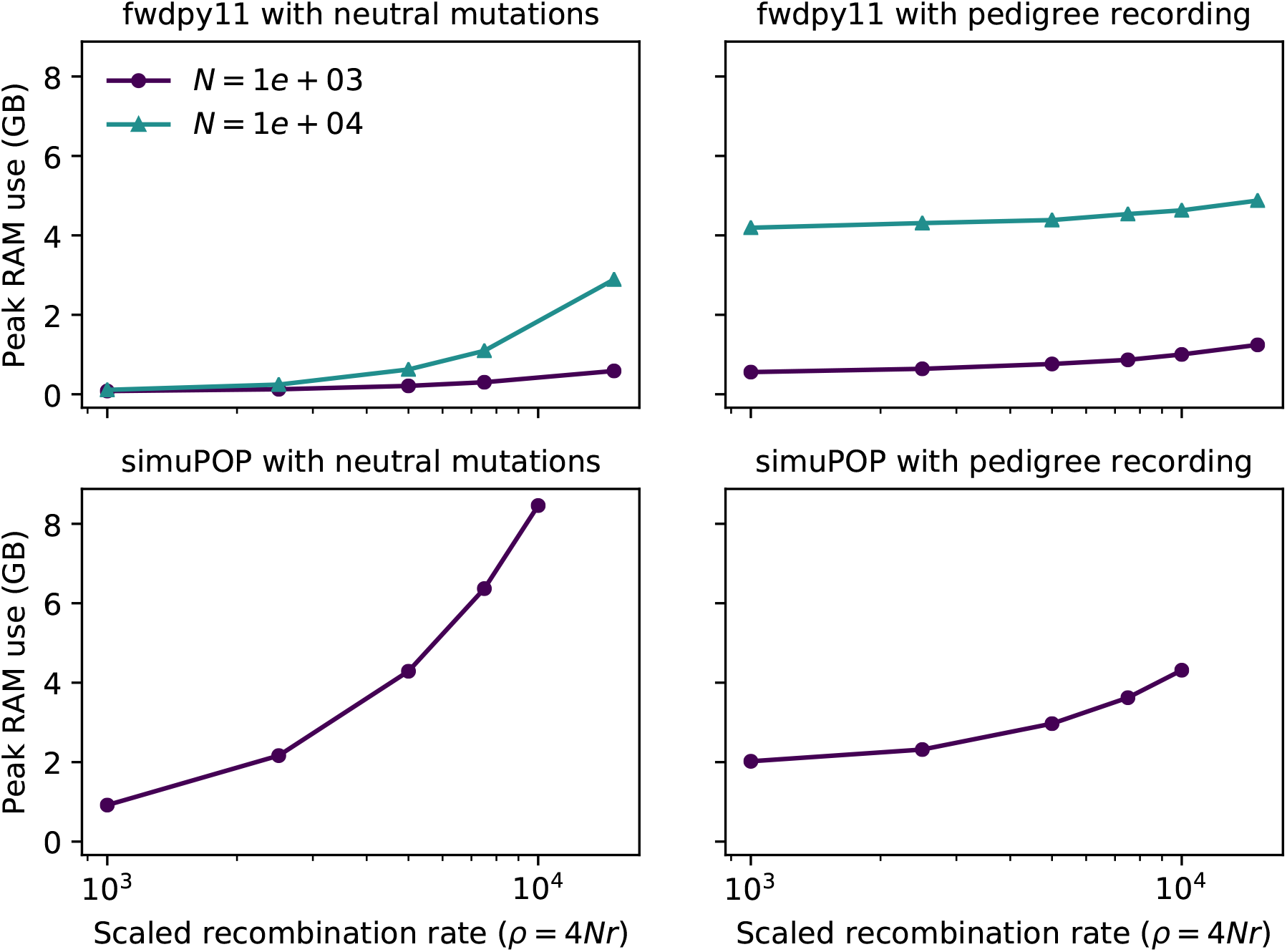
Peak RAM use in gigabytes (GB) for simulations without selection. The plots are from the same data as in Figure D1.

**Figure B1:**
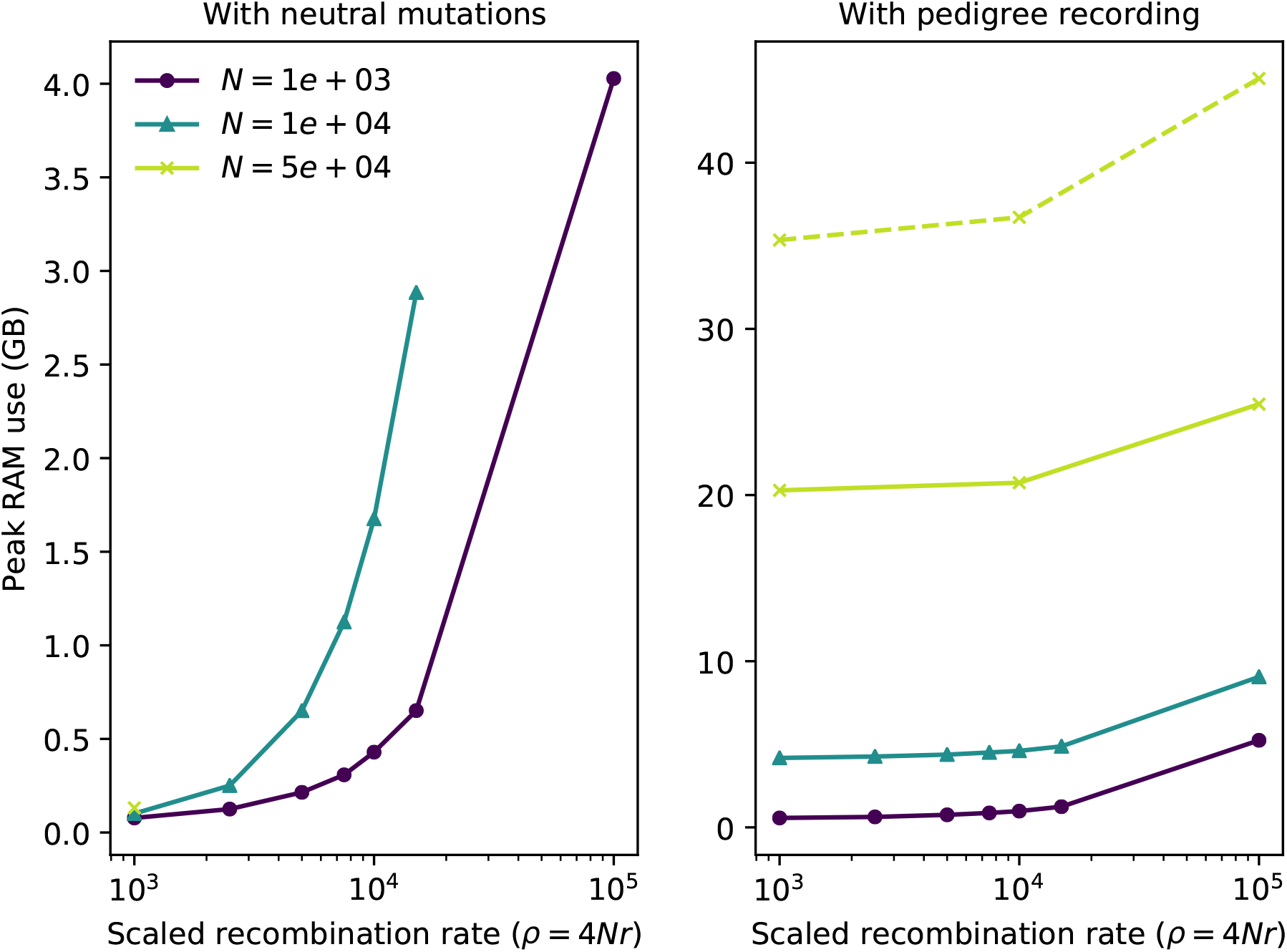
Peak RAM use in gigabytes (GB) for simulations with selection. The plots are from the same data as in Figure 1. Note that the y-axis scales differ considerably across panels.

## B Memory usage with selection

The peak RAM use in gigabytes (GB) is shown for the simulations with selection in Figure B1. For our parameters, which involved simplifying every 10^3^ generations, simulation while recording pedigrees uses more RAM than a standard simulation. This is because with pedigree recording, a non-recombinant individual still requires one new edge and two new nodes to be allocated. Without pedigree recording, no new memory is allocated for such an offspring by `fwdpp`. Thus, pedigree recording accumulates a large amount of data in RAM until simplification. This extra RAM consumption is the trade off for reduced run times and the degree of extra RAM needed can be reduced by simplifying more often. The simulations where we run the simplification step in a separate thread require even more RAM (dashed line in Figure B1) because up to four sets of unsimplified data are stored in RAM. In practice, the RAM consumption can be minimized by simplifying more often.

## C Effect of simplification interval on run time and memory use

We performed a limited set of simulations to explore the effect of the simplification interval on run times and memory use. Figure C2 shows that run times increase for simplification intervals less than 100 generations. For intervals longer than 100 generations, run times are very similar, with no appreciable difference between 10^3^ and 10^4^ generations. As expected, the simplification interval has a near-linear effect on peak RAM use (Figure C3). Taken together Figures C2 and C3 suggest that one can tune the simplification interval to available RAM with little effect on run times provided that simplification doesn’t occur too often.

**Figure C2:**
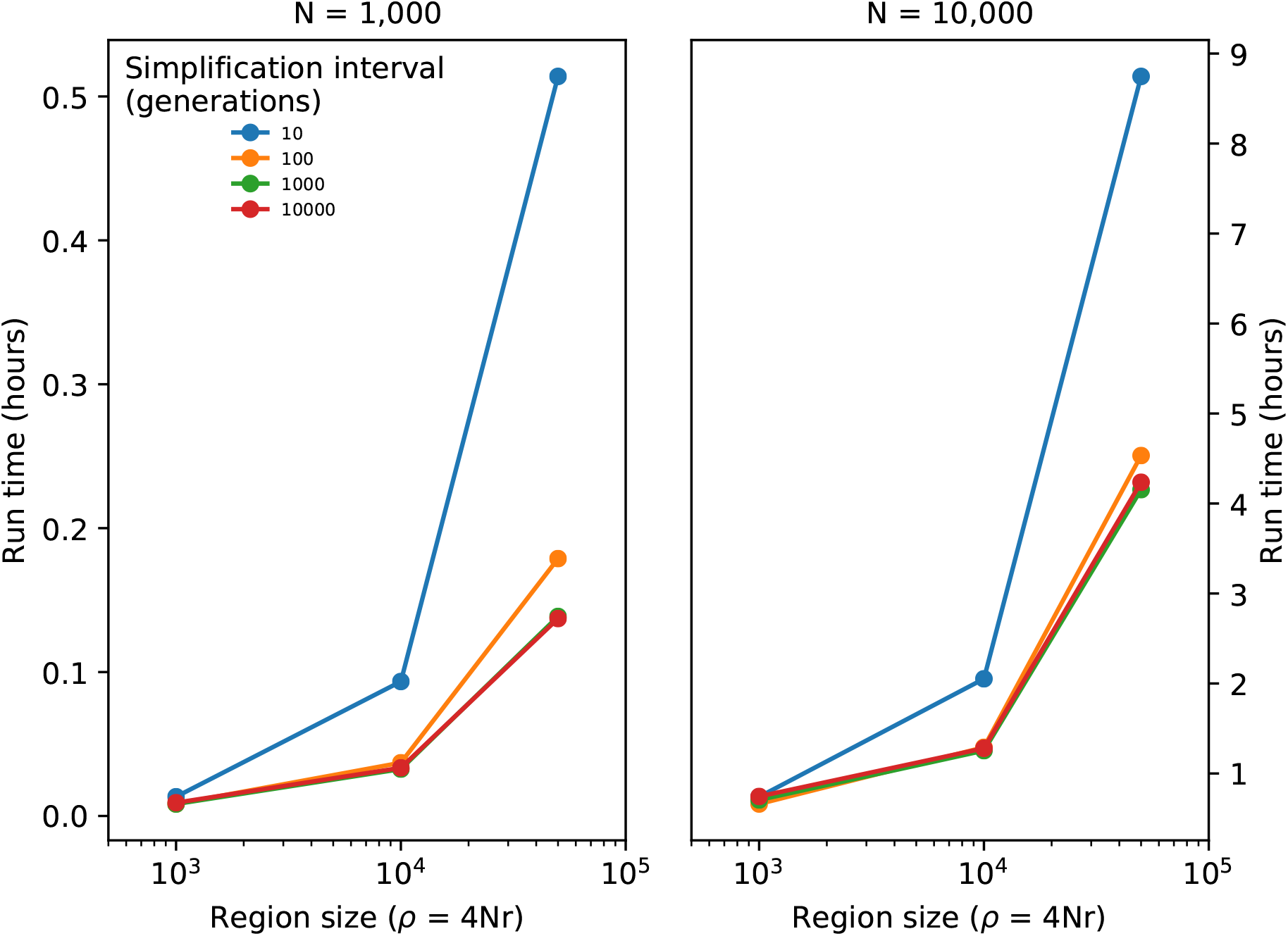
The effect of simplification interval on run times. A single replicate was done for the parameters shown in the figure. These simulations were done with the same selection parameters as in Figure 1.

**Figure C3:**
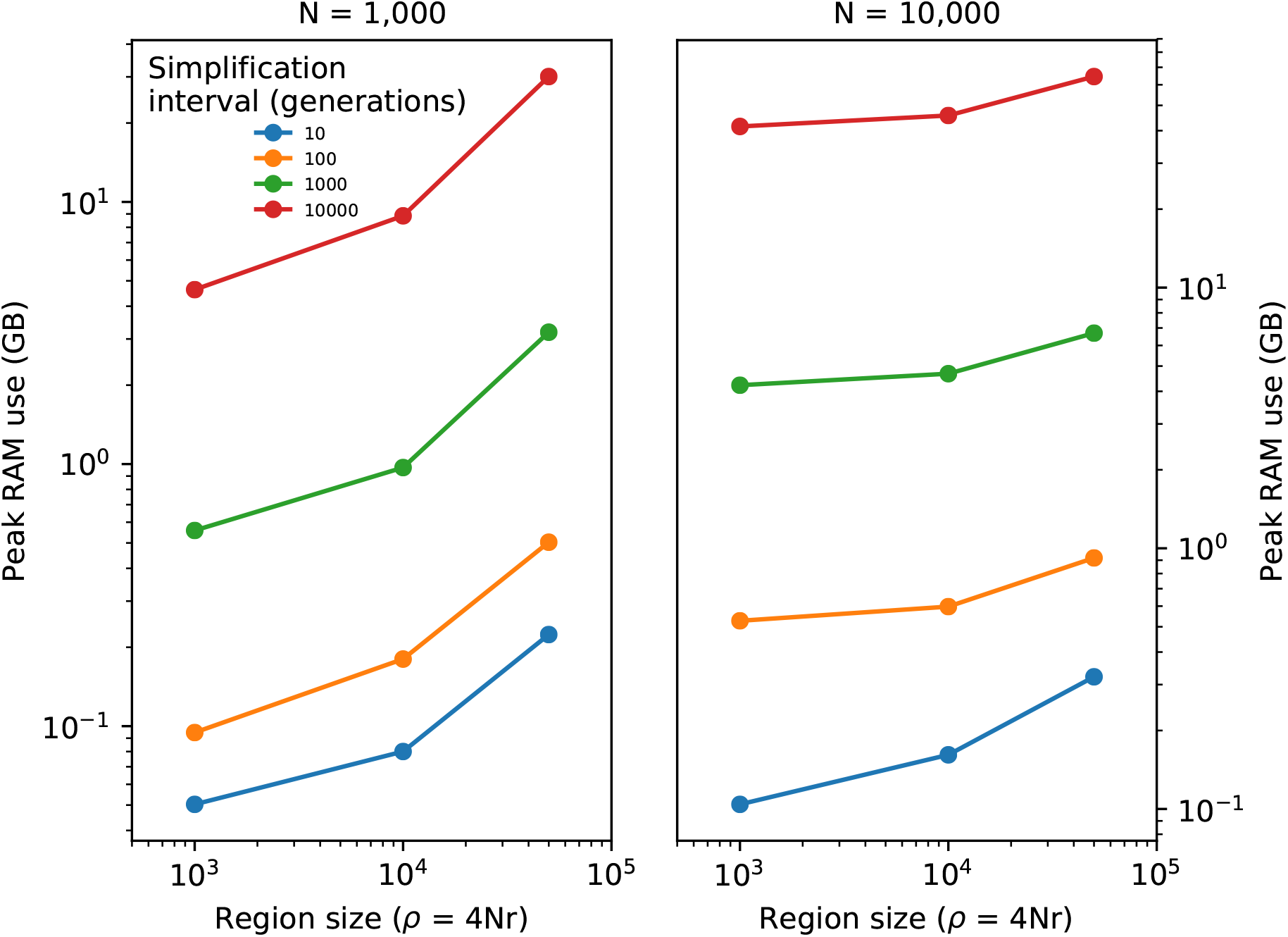
Peak RAM use for the simulations shown in Figure C2.

## D The simuPOP implementation

To use the methods with simuPOP, we created a python module (available at https://github.com/ashander/ftprime) that manages communication between the forward simulator and `tskit`. We implemented simulations in a Python script, usable as a command-line utility, taking input parameters for population size *N* and scaled recombination rate *ρ*. As for the fwdpp simulations above, we set scaled mutation rate *θ* = 4*Nμ* = *ρ* (where *μ* is the per-site mutation rate multiplied by the number of sites), and used a simplify interval of 1000 generations and simulated for *T* = 10*N* generations in each simulation.

We coupled `simuPOP` to `tskit` using a Python class `RecombCollector` that 1) tracks time, and 2) collects and parses recombination information provided by the `Recombinator` class of `simuPOP`, adding them to a **NodeTable**() and **EdgeTable**() upon which simplification is performed every 1000 generations (using `tskit.sort_tables`() and `tskit.simplify_tables`). The tables are initialized using the `TreeSequence` returned from `tskit.simulate`(2*N*, `Ne=`*N*, `recombinationjrate=`*r*/2, `length=`*n*). The `simuPOP.Population`() is initialized and tagged with individual IDs, a mapping of these IDs to the **NodeTable**() IDs is passed to an instance of `RecombCollector` and used internally to update the tables. The `simuPOP.Population`() is then evolved. Before mating: the internal time of the `RecombCollector` instance is updated. During mating: male and female parents are drawn randomly, recombination data is output by `simuPOP` to the `RecombCollector` instance. After mating: every 1000 generations the tables are simplified.

`RecombCollector` uses the Python class `ARGrecorder` internally to add data to the tables and perform simplification. This class contains the **NodeTable**() and **EdgeTable**() and also the mapping from (haploid) ids to ids in the tables. It also provides methods (`add_individual` and `add_record`) to append data to the tables. The only functions of `RecombCollector` not delegated to this internal class are computing haploid IDs from `simuPOP`’s diploid IDs and parsing recombination data into data suitable for use with use with `add_individual` and `add_record`; these functions are used to stream data into the tables as it is received by an instance of `RecombCollector`.

## E Simplification algorithm

Here we provide a concrete implementation of the simplification algorithm. The algorithm uses a few simple structures besides the node and edge tables discussed earlier. As the algorithm needs to track the mapping between input and output node IDs over specific genomic intervals, we use a Segment () type to track these individual mappings. Thus, a segment *x* has three attributes: *x*.left and *x*.right represent the genomic interval in the usual way, and *x*.node represents the output node ID that is assigned to this interval. These output node IDs correspond to colors in the “paint pot” analogy. The per-interval segments are assigned to input IDs via the 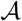 mapping, which is the main state required by the algorithm. Each element 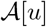 is a list of ancestral segments, describing the output node IDs mapped to input node ID *u* over a set of intervals. For each input node, we need to process the ancestral segments to find overlaps (and hence coalescences), and to update 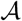. We do this using a **PriorityQueue**() type, which maintains a list of ancestral segments sorted by left coordinate. For a priority queue *Q*, the operation *Q*.**push**(*x*) inserts the segment *x* into the queue, and *Q*.**pop**() removes and returns the segment in *Q* with the smallest left coordinate. To obtain the segment currently in the queue with the smallest left value we write *Q*[0]. Finally, note that to keep the exposition simple, the algorithm described here requires that the input node table is sorted by time since the parent’s birth; but the version implemented in tskit makes different sortedness assumptions (described in the documentation).

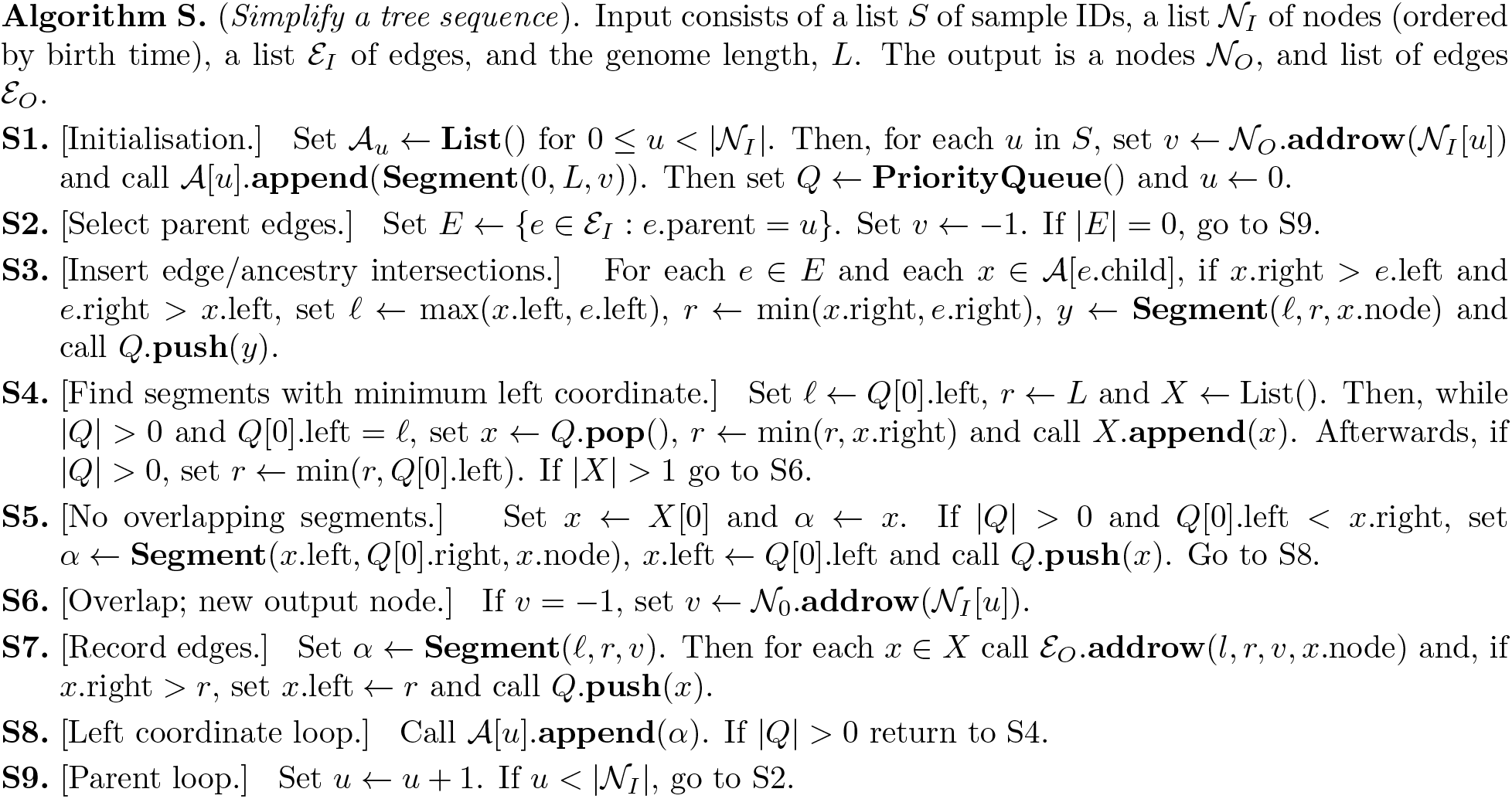

The algorithm begins in S1 by allocating an empty list to store ancestral segments for each of the 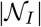 input nodes, and then creating the initial state for the samples in *S*. For each input sample with node ID *u*, we add a row in the output node table 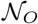, and create a mapping for this new node *v*. After this initial state has been created, we then allocate our priority queue *Q*, and set the current input ID *u* to 0. Recall that the function 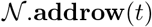 adds a new node to the node table 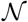 with birth time *t*, and returns the ID (i.e., index) of that new node in 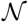. So, the operation 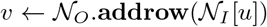 looks up the birth time for input node ID *u* in 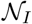, adds a new node with this time to 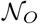, and stores the resulting (output) node ID in *v*.

To process ancestral information moving back up through the pedigree, steps S2–S9 consider each node u in turn (note we are assuming that nodes are ordered by time-since-birth). In S2 we first find all edges in the input where the parent is equal to our current input ID *u*, and then set the corresponding output ID *v* to −1. Then, in S3 we find all intersections between these input edges and the mapped ancestry segments of their child nodes. For each of these intersections we create a new segment *y* and insert it into the priority queue. (In the paint pot analogy, this stores in *Q* all segments of color that must be transferred to the ancestor with input ID *u*.)

After finding all overlaps and filling the priority queue with the resulting ancestry segments, we then need to process these segments. Steps S4-S8 loop over the segments in the queue, considering each in turn until the queue is empty. Step S4 builds a list *X* of all segments covering the current left-most coordinate (along with some bookkeeping to keep track of where our next right coordinate *r* is), removing them from *Q*. If there is only one element in *X* this means that we have no overlapping ancestral segments at this coordinate and we proceed to S5.

In this case, only a single segment covers the leftmost region of the segments, so there is no change in the mapping between input and output IDs, and the ancestry segment α is not changed (this is assigned to 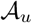 in S8). (The color in this segment of the offspring can simply be transferred to the parent.) However, if other segments in *Q* intersect with *x* (since *x* is the current left-most segment, these have left coordinate less then *x*.right), then we can only transfer the portion of *x* up until it overlaps with other segments. Therefore, we change the left coordinate of *x* to this coordinate and reinsert it into *Q* where it will be processed again later, effectively cutting off (and propagating) the leftmost segment of *x*. After doing this, we proceed to S8, update the ancestry mapping for the input node *u*, and loop back to S4 if *Q* is not empty.

On the other hand, if there is more than one segment in *X*, we know that an overlap among the ancestral segments has occurred and we must update the output nodes and edges accordingly. In step S6 we first allocate a new output node *v* for input node *u*, if it has not been done already (we may have many overlapping segments along the genome). Step S7 then continues by creating a new ancestry segment *α* mapping the current interval to the output node *v*. We then iterate over all segments in *X*, adding the appropriate output edges and editing and reinserting the ancestry segment *x* into the priority queue if necessary, as before.

## Python implementation of simplify (supplementary information)

”””

*Python implementation of Algorithm S*.

*The example works by generating an initial TreeSequence for a sample of 10 haplotyp es using msprime. We then simplify the node/edge table in that TreeSequence with respect to the first three samples*.

**import** heapq

**import** numpy as np

**import** msprime

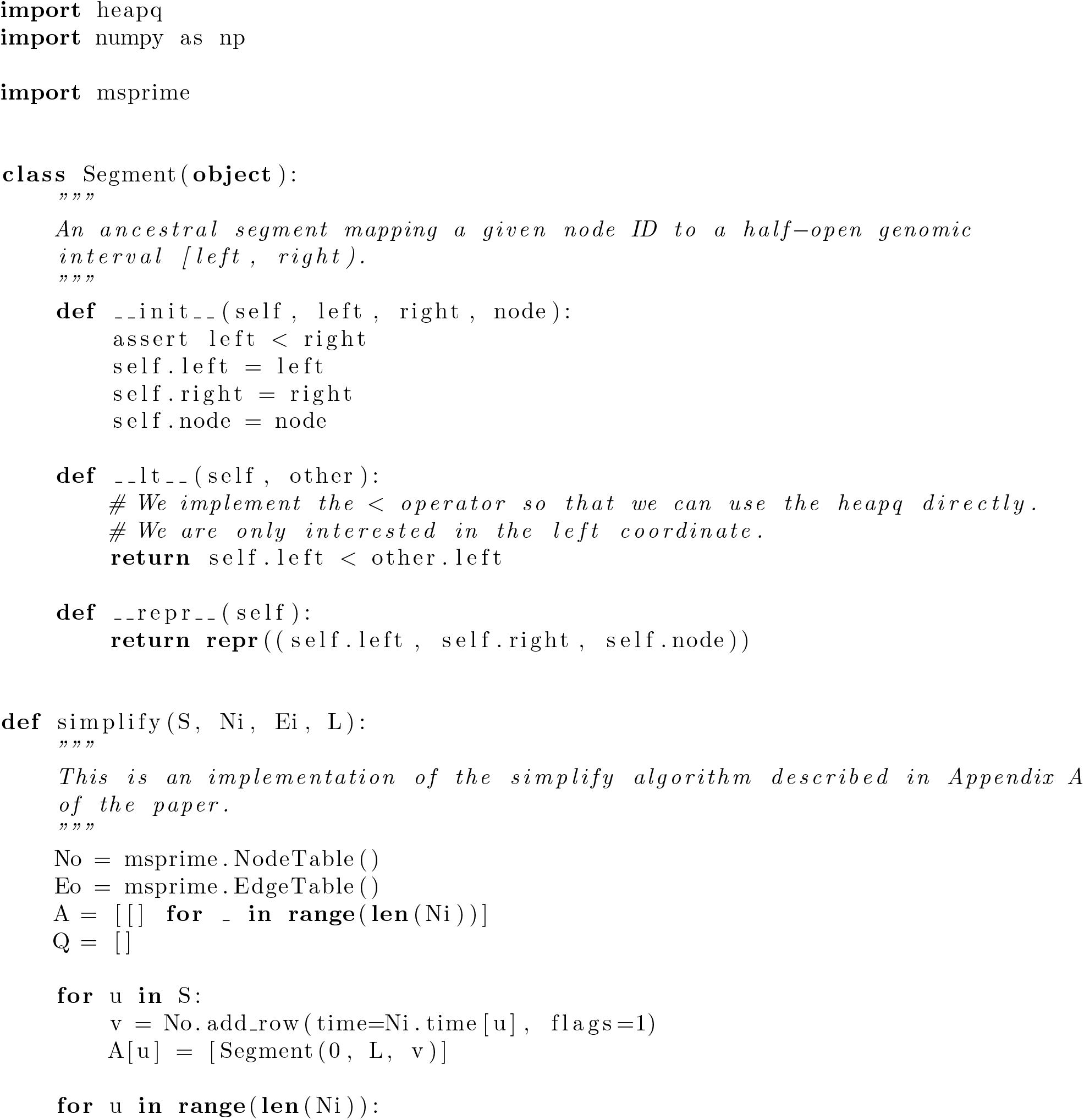
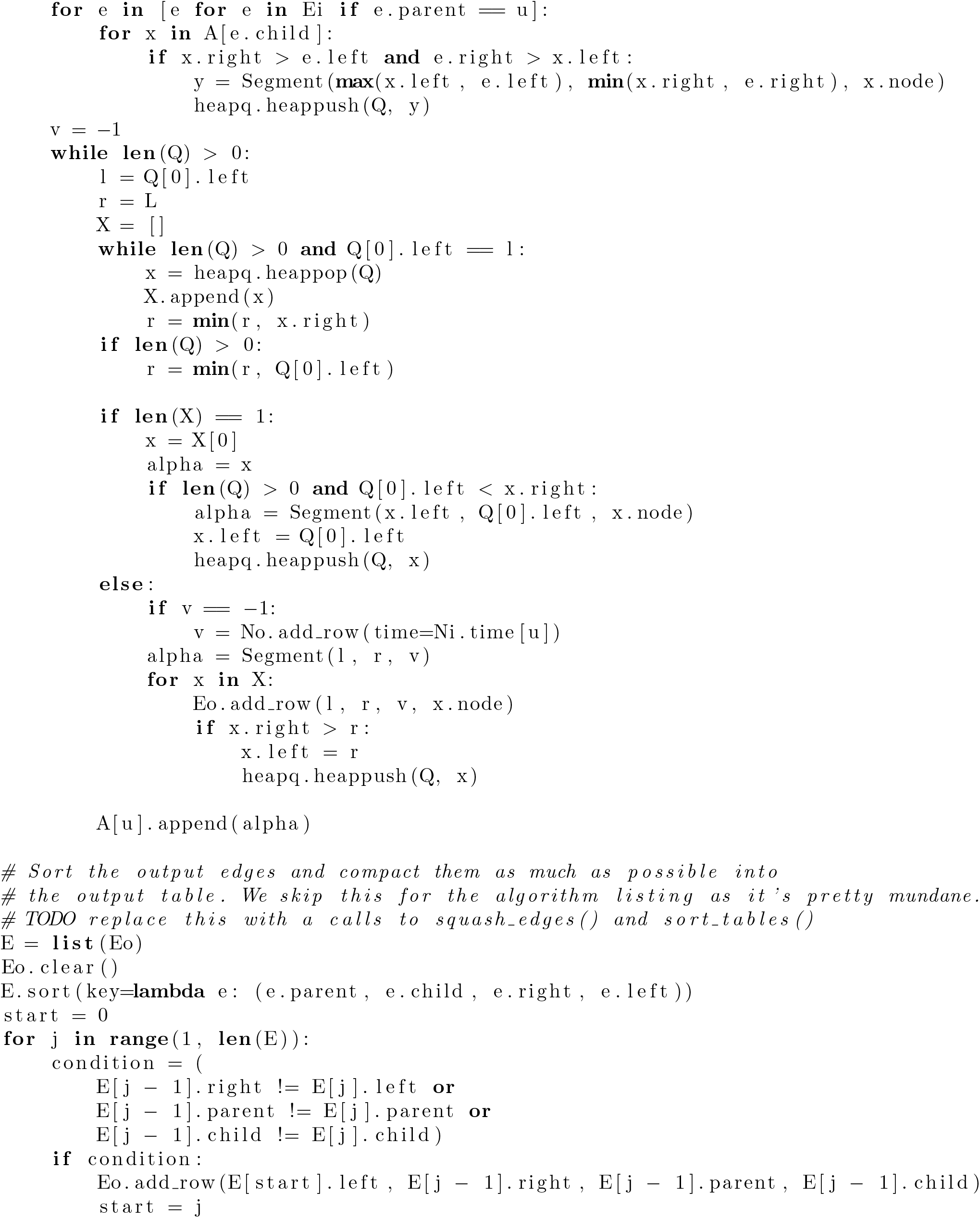
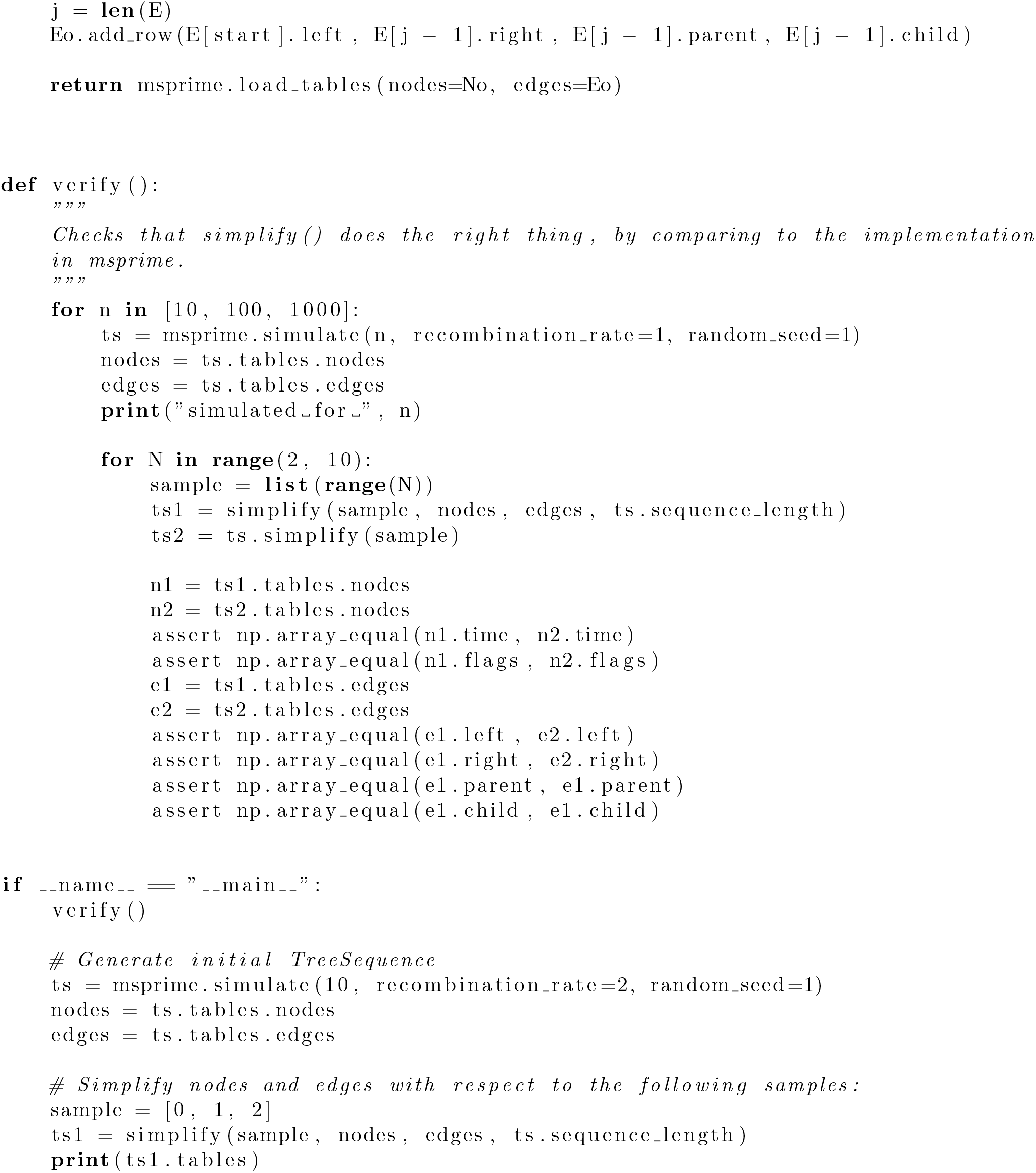

